# Viral infection of algal blooms leaves a halogenated footprint on the dissolved organic matter in the ocean

**DOI:** 10.1101/2020.09.08.287805

**Authors:** Constanze Kuhlisch, Guy Schleyer, Nir Shahaf, Flora Vincent, Daniella Schatz, Assaf Vardi

## Abstract

Algal blooms are important hotspots of primary production in the ocean, forming the basis of the marine food web and fueling the pool of dissolved organic matter (DOM)^1^, which is the largest global inventory of reduced carbon and a market place for metabolic exchange in the ocean^2^. Marine viruses are key players in controlling algal bloom demise and act as major biogeochemical drivers of nutrient cycling and metabolic fluxes by shunting algal biomass from higher trophic levels to the DOM pool, a process termed the ‘viral shunt’^3,4^. Nevertheless, the metabolic composition of virus-induced DOM (vDOM) in the marine environment is unknown. To decode the metabolic footprint of the ‘viral shunt’, we induced a bloom of the ecologically important alga *Emiliania huxleyi* in the natural environment, and followed its succession using an untargeted exometabolomics approach. Here we show that algal bloom succession induces extensive and dynamic changes in the exometabolic landscape, especially during bloom demise. By correlating to a specific viral gene marker, we discovered a set of novel chlorine-iodine-containing metabolites that were induced by viral infection and copiously released during bloom demise. We further detected several of these chloro-iodo metabolites in virus-infected open ocean blooms of *E. huxleyi*, supporting their use as sensitive biomarkers for virus-induced demise in the natural environment. Therefore, we propose halogenation to be a hallmark of the *E. huxleyi* vDOM, providing insights into the profound metabolic consequences of viral infection for the marine DOM pool.

## Main text

Marine dissolved organic matter (DOM), estimated at 662×10^12^ kg, is the largest inventory of reduced carbon in the ocean^2^. It is composed of more than 100,000 different molecules^5^, of which over 95% are stable (recalcitrant) for thousands of years^6^. Photosynthetic organisms are the main source of DOM in the sunlit ocean, particularly of bioavailable (labile) molecules that are rapidly turned over by the microbial community^7-9^. Algal blooms, events of increased phytoplankton biomass, significantly contribute to the release of labile metabolites^10^. However, it is largely unknown how algal bloom succession shapes DOM composition and how different biotic interactions impact the DOM pool. A major driver of bloom demise is lytic viral infection, which leads to the release of algal biomass to the DOM pool, a process termed the ‘viral shunt’^3,4^. This key ecosystem process links primary production with the DOM pool and curtails the energy transfer to higher trophic levels, hence affecting the biogeochemical cycling of major nutrients, such as carbon, nitrogen and sulfur, and fueling the microbial food web in the ocean. Laboratory-based experimental approaches suggest that this virus-induced DOM (vDOM) has a characteristic composition^11^. However, the chemical nature of vDOM, driven by the ‘viral shunt’ during algal bloom demise, has not yet been explored in the natural environment. Moreover, the characterization of individual metabolites that are unique to the vDOM is still lacking. An ecologically important host-virus model system is the cosmopolitan alga *Emiliania huxleyi* and its specific large dsDNA virus, *E. huxleyi* virus (EhV). *E. huxleyi* frequently forms vast blooms in the ocean, which are routinely terminated by lytic viral infection^12-14^. These blooms are an important biomass source for the marine food web and affect the global biogeochemical cycling of carbon and sulfur^15-19^. Viral infection leads to profound remodeling of the *E. huxleyi* metabolic pathways to support infection, including enhanced glycolytic fluxes, elevated fatty acid synthesis and production of specific virus-derived glycosphingolipids^20-23^. Once cells lyse, the metabolism of infected cells can act as a source for DOM, releasing a unique bouquet of metabolites to the ocean. In this study, we aimed to characterize the DOM composition during the succession of an induced phytoplankton community by applying an untargeted exometabolomics approach. This led to the discovery of novel halogenated metabolites that are hallmarks of virus-induced bloom demise of *E. huxleyi* in the natural environment.

Phytoplankton bloom succession shapes the marine DOM We sought to examine the impact of phytoplankton bloom succession, in particular of *E. huxleyi*, on the composition of marine DOM by using an *in situ* mesocosm setup in the coastal waters of southern Norway (Extended Data Fig. 1a), where annual blooms and viral infection of *E. huxleyi* occur naturally. Natural marine microbial communities were enclosed in four mesocosm bags and monitored daily over 24 days (Extended Data Fig. 1b). All bags were supplemented with nutrients at a nitrogen to phosphorous ratio of 16:1 to favor the growth and induce a bloom of *E. huxleyi*^24^. The phytoplankton community in each bag responded with an increase in biomass compared to the surrounding fjord, as indicated by elevated chlorophyll levels (Fig. 1a). A first bloom of a mixed algal community peaked at day 10, consisting mainly of pico- and nanoeukaryotes (Extended Data Fig. 2). This was followed by a bloom of *E. huxleyi*, as indicated by an increase in calcified cells, reaching up to 8×10^7^ cells/L (Fig. 1b).

**Fig. 1:**
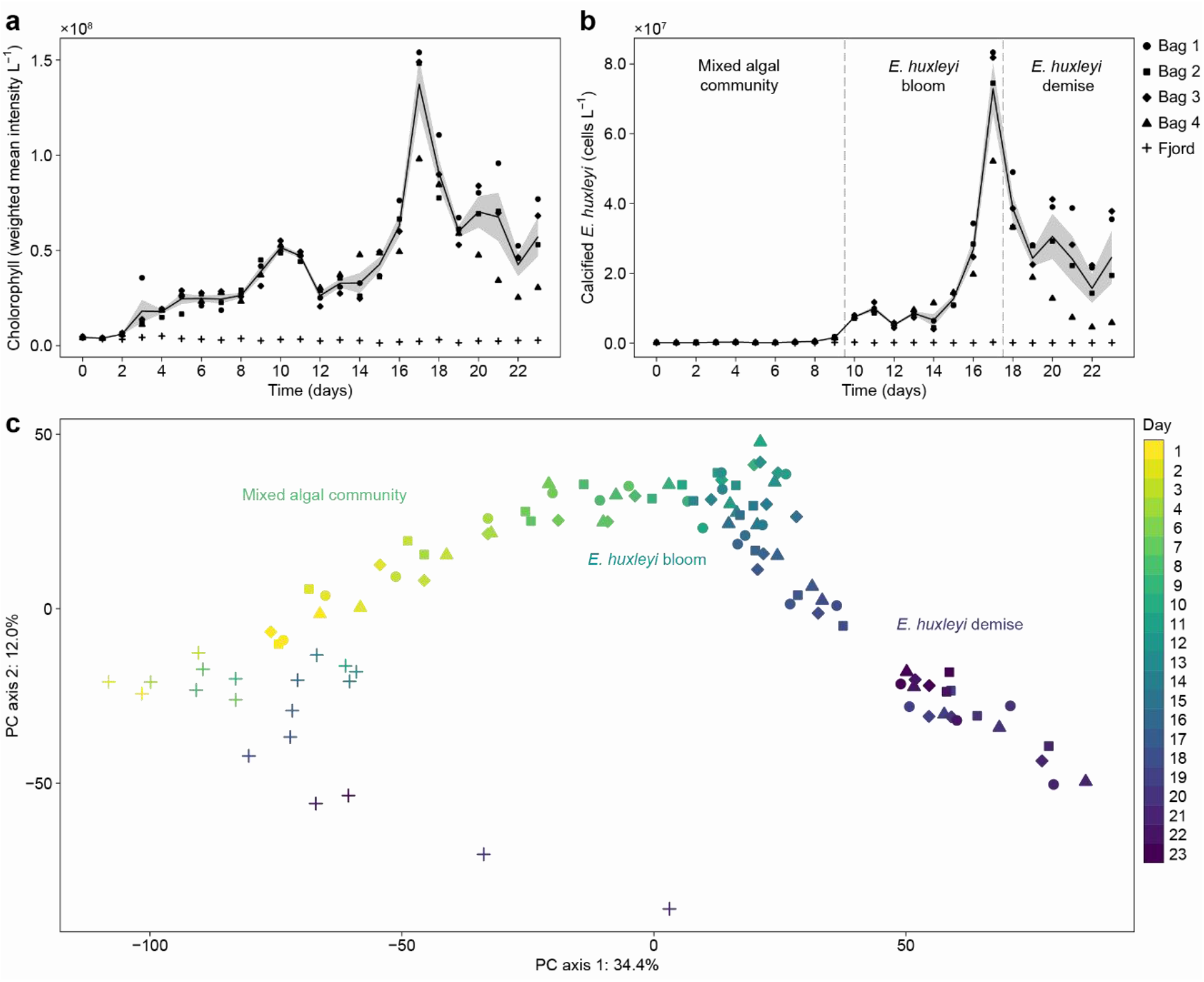
Phytoplankton bloom succession is a major driver of marine DOM composition. **a**, Induced phytoplankton growth and succession in four mesocosm bags (filled symbols) compared to the surrounding fjord (crosses) following nutrient amendments. **b**, Growth of the calcifying alga *E. huxleyi* revealed three distinct phases: an initial bloom dominated by a mixed algal community, the bloom of *E. huxleyi* and its demise (separated by dashed lines). Chlorophyll levels (a) and calcified cell abundance (b) are based on flow cytometry analysis. For both, the average (solid line) and standard error (grey shading) of all bags (n = 4) are indicated. **c**, Principal component analysis (PCA) separated the exometabolite profiles of fjord (crosses) from mesocosm bag (filled symbols, as in a and b) samples, with PC axis 1 reflecting bloom succession through time, revealing its impact on DOM composition. An untargeted metabolomics approach (revealing 6,787 mass features) was used for exometabolite profiling. Experimental days are indicated by symbol color, from yellow (day 1) to blue (day 23).

Based on *E. huxleyi* cell abundance, we defined three phases: the mixed algal community phase with a low abundance of calcified cells (<1×10^6^ cells/L; days 0-9), the bloom phase with an increase in calcified cells (days 10-17), and the demise phase, showing a sharp decrease in calcified cells (days 18-23). Phytoplankton bloom succession occurred similarly in all four bags apart from the demise phase of *E. huxleyi*, during which the bags diverged from one another (Fig. 1a, b).

To characterize the composition and monitor changes in the DOM pool that are derived from biotic interactions during bloom succession, we applied an untargeted exometabolomics approach, in which we profiled small, semi-polar, extracellular metabolites (Extended Data Fig. 1c). Seawater filtrates were extracted daily using solid phase extraction (SPE) cartridges, and their metabolite composition was analyzed by liquid chromatography-high resolution mass spectrometry (LC-HRMS). Principal component analysis (PCA) of the extracted exometabolites (6,787 mass features) revealed a clear separation of fjord water samples from bloom-associated mesocosm samples throughout the experiment (Fig. 1c). The exometabolomes of the mesocosm bags were separated along the first PC axis (34.4%), which likely represents phytoplankton community succession over time. Furthermore, changes in metabolite composition between days were more pronounced than the variability between bags (repeated-measures ANOVA, *P* <0.0001). Interestingly, the exometabolomes changed more substantially during the mixed algal community phase and the demise phase of *E. huxleyi* compared to a relatively stable exometabolic landscape during the *E. huxleyi* bloom phase (Extended Data Fig. 3).

We further investigated the DOM composition at the metabolite level to reveal the effect of bloom succession on the production and release of exometabolites as opposed to their consumption or degradation. About 26% (n = 1,747) of all detected mass features were elevated during succession of the induced phytoplankton community compared to the fjord. Hierarchical cluster analysis (Fig. 2a) revealed two distinct occurrence patterns: mass features that were abundant during the mixed algal community phase (n = 518), and mass features that increased during the *E. huxleyi* bloom and demise phase (n = 1,229). Closer inspection of six selected clusters (Fig. 2b) revealed diverse dynamics and fluctuation patterns in the bloom exometabolome.

**Fig. 2:**
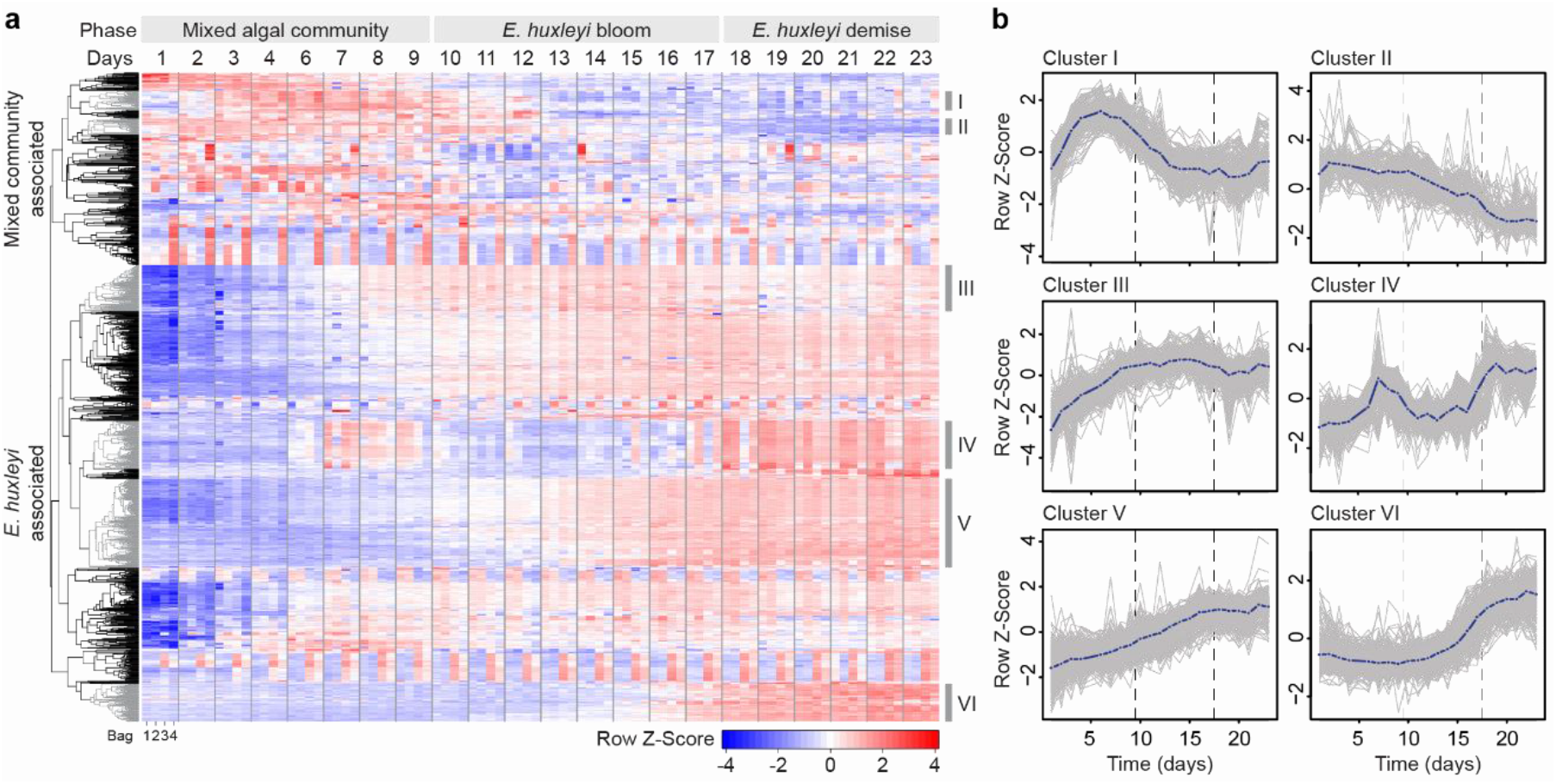
Dynamic changes in the exometabolic landscape during algal bloom succession. **a**, Hierarchical cluster analysis and heat map of exometabolites that increased following the induction of phytoplankton blooms in mesocosm bags compared to the fjord (1,747 mass features). Samples (columns) are ordered by time along bloom succession (days), each day showing all four bags (bag 1-4). Mass features (rows) are hierarchically clustered based on their log-transformed and standardized intensity profiles. Algal bloom phases are indicated on top. Six clusters with notable temporal profiles are highlighted in grey (Cluster I-VI). **b**, Temporal profiles of the selected clusters reveal diverse patterns driven by biological processes. Individual intensity profiles (grey lines) are shown together with an average cluster profile (blue line). Algal bloom phases are separated by dashed lines.

Some metabolites increased rapidly within a few days (Cluster I, VI), whereas others increased gradually over more than two weeks (Cluster V), indicating several biogenic sources throughout bloom succession. Metabolite consumption or transformation either occurred within a few days (Cluster I), steadily over several weeks (Cluster II), or were not observed until the end of the experiment (Cluster III), thus representing a range of exometabolites with different lability and turnover times. Interestingly, some metabolites peaked during both the mixed algal community and the *E. huxleyi* bloom phase (Cluster IV), suggesting a general association with phytoplankton blooms.

A major driver of the exometabolite composition was the phytoplankton bloom succession as shown by chlorophyll levels (Pearson correlation, *r* = 0.62), compared to abiotic conditions such as temperature (Pearson correlation, *r* = 0.03). Accordingly, the fjord water exometabolome showed only minor changes throughout the experiment (Fig. 1c). Phytoplankton bloom succession was previously shown to affect the recalcitrant DOM^25^. Here, facilitated by the high temporal resolution of our metabolite profiling, we show the important role of bloom succession in shaping the biologically active, labile DOM. The metabolite profiles further indicate that *E. huxleyi* blooms stabilize the chemodiversity of marine DOM. Thus, the extent to which blooms orchestrate DOM composition seems to depend on the characteristics of the bloom, such as species dominance. Some of the major oceanic phytoplankton blooms may hence be characterized by distinct metabolite footprints, which could drive the growth of specialized associated microbial communities^26^. In comparison, both the mixed algal community phase and the *E. huxleyi* demise phase showed strong changes in the exometabolome. This might be explained by a fast succession of primary producers competing for the newly available inorganic nutrients during the mixed algal community phase, and by a fast succession of microbial consumers competing for the released organic matter during the *E. huxleyi* demise phase. By applying an untargeted exometabolomics approach daily over several weeks, we revealed the highly dynamic changes in the exometabolic landscape that occur throughout phytoplankton bloom succession. The majority of metabolites that are released during algal blooms are rapidly transformed or consumed by microbes within hours or days, hampering their discovery^6^. Here, we were able to shed light on this biologically active, labile DOM fraction, thus expanding our ability to track changes in the DOM composition. Although recent developments in analytical chemistry significantly improved the ability to describe DOM composition, major challenges remain to decode the richness of seawater chemodiversity and to unambiguously identify its components. Another major bottleneck is to link changes in DOM composition to specific microbial interactions that occur during microbial community succession. Of specific interest are host-virus interactions, which can lead to profound remodeling of host metabolism and to lysis of host cells, thereby having major biogeochemical consequences^27,28^. However, the consequences of virus-induced bloom demise for DOM composition are poorly understood.

### The metabolic footprint of alga-virus interaction

We aimed to discover exometabolites that are specific to virus-induced *E. huxleyi* bloom demise. Infection dynamics of *E. huxleyi* varied among the bags (Fig. 3a), providing a comparative tool to identify metabolites that are highly enriched during viral infection. The strongest increase in viral production occurred in bag 4, followed by bag 2 and bag 1, whereas in bag 3 no viral proliferation was observed. This was also reflected in *E. huxleyi* bloom demise, which varied between the bags, reaching the lowest cell abundance in bag 4 (Fig. 3a). To resolve the metabolic composition of the virus-induced DOM (vDOM), we correlated the temporal profile of each metabolite with a gene marker of EhV (major capsid protein, *mcp*). In total, 705 mass features were positively correlated with *mcp* abundance during the bloom and demise phase in the most infected bag 4 (Pearson correlation, *r* >0.7, p < 0.05). Hierarchical cluster analysis (Extended Data Fig. 4) and inspection of the intensity profiles from all bags revealed a subset of 65 mass features that were differential between the bags and corresponded to the different levels of viral infection. Following feature deconvolution and manual curation, these mass features were grouped into 20 metabolites (Fig. 3b; Extended Data Table 1). This metabolic repertoire characterizes the vDOM produced by virus-induced bloom demise of *E. huxleyi*.

**Fig. 3:**
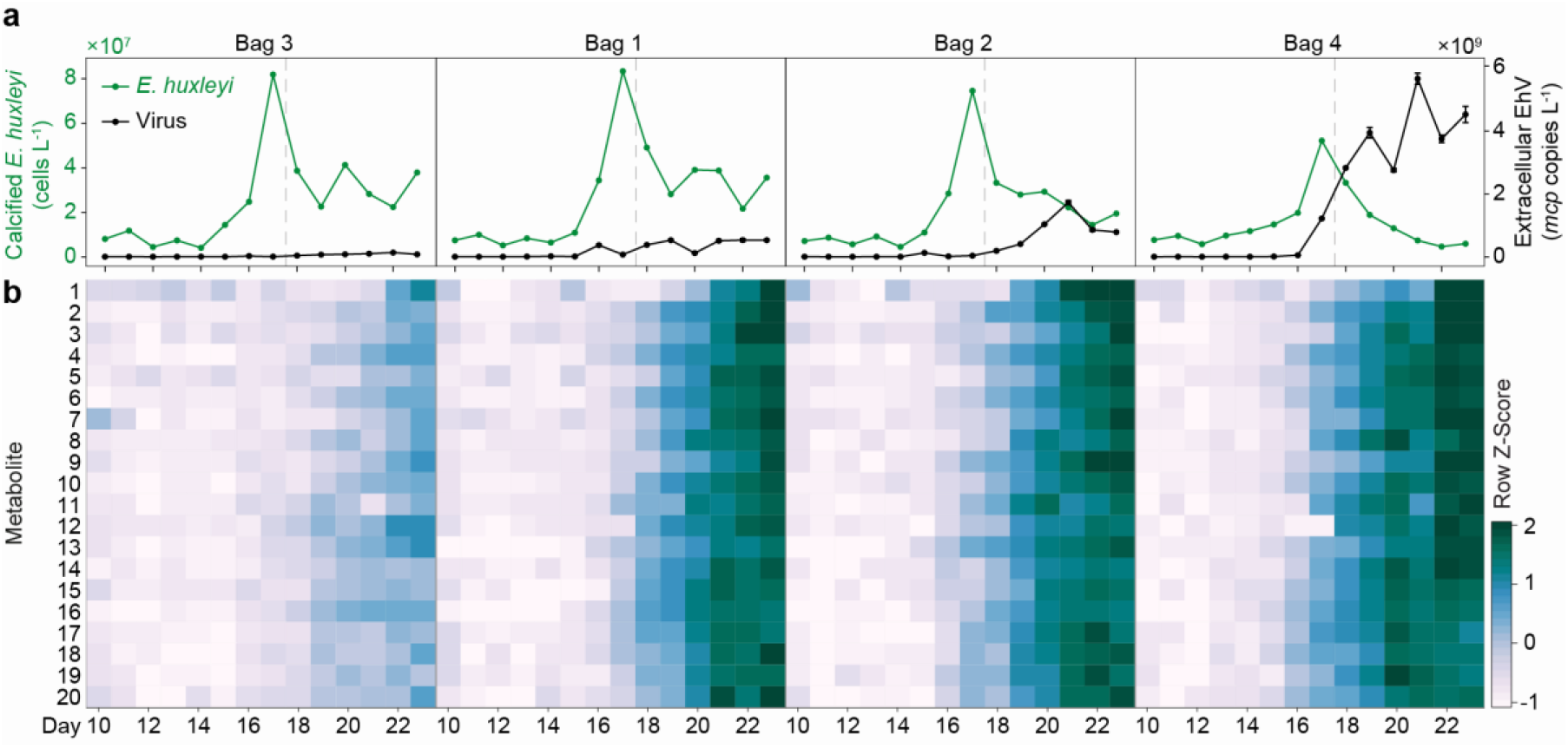
Lytic viral infection of *E. huxleyi* blooms leaves a distinct metabolic imprint on the marine DOM pool. **a**, Variable host-virus dynamics across the mesocosm bags based on the abundance of *E. huxleyi* cells (green) and extracellular EhV (black). Bags are ordered by EhV abundance. All *E. huxleyi* blooms peaked at day 17, with the highest cell abundance reached in bag 3 and the lowest in bag 4. The onset of bloom demise is indicated (dashed line). Major capsid protein (*mcp*) copy values are presented as average ± standard deviation (n = 3). **b**, Heat map of 20 metabolites that showed an intensity increase in response to viral infection. For each metabolite, the intensity of the most abundant mass feature ([M+H]^+^ or [M+H-H2O]^+^) is shown after log-transformation and standardization. Metabolites are ordered according to retention time (Extended Data Table 1).

Intriguingly, structural characterization of the newly identified vDOM metabolites revealed that 17 out of 20 are halogenated, having different combinations of chlorine, bromine and iodine (Extended Data Table 1). Specifically, nine vDOM metabolites contained 2-3 chlorine atoms as well as iodine based on the isotope distribution and MS/MS analysis (Fig. 4a, b, Supplementary Data). The predicted molecular formulas (Extended Data Table 1) were not found in common mass spectral and natural product libraries.

**Fig. 4:**
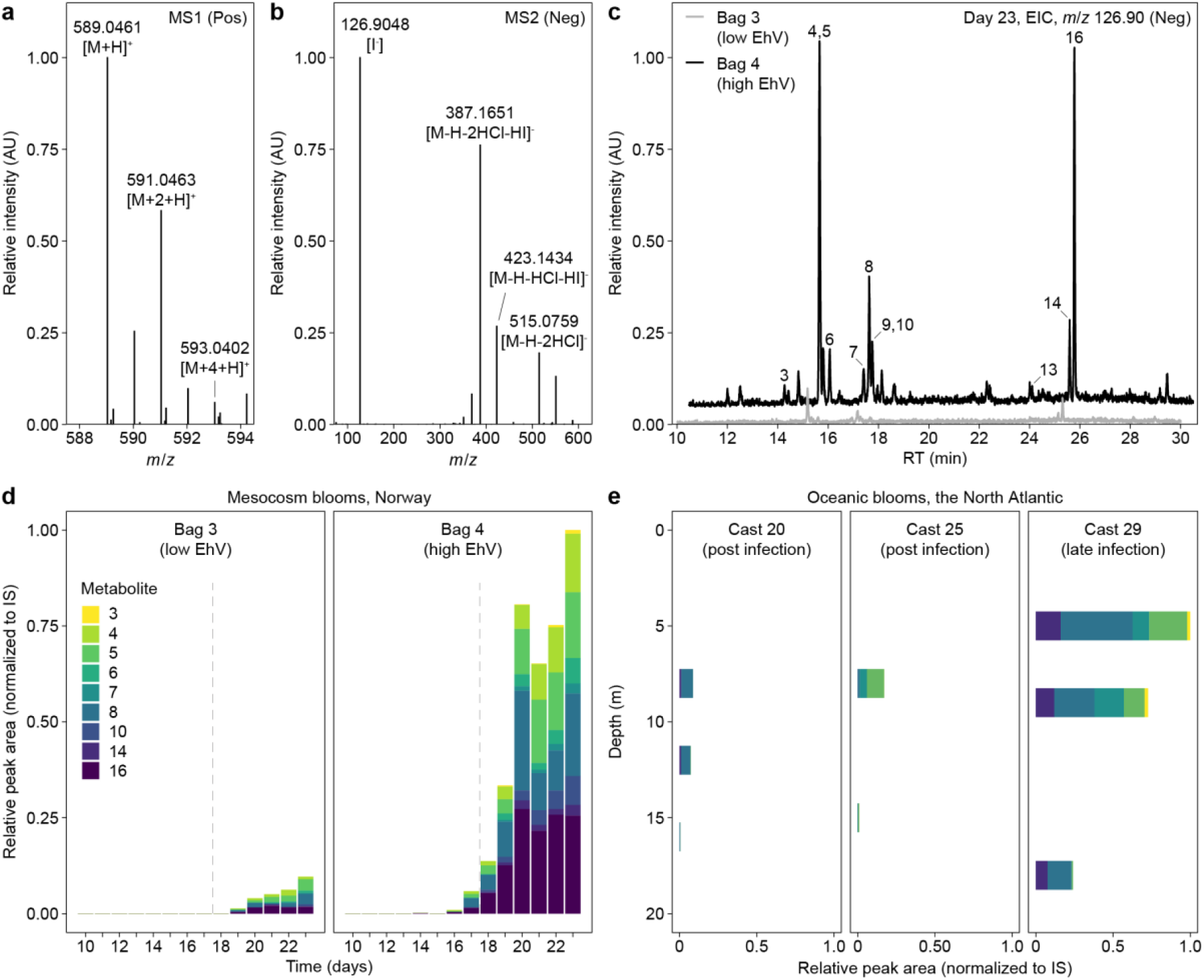
The vDOM produced during *E. huxleyi* bloom demise is characterized by unique chlorine-iodine-containing metabolites. **a**, Mass spectrum of the molecular ion with *m*/*z* 589.0464, [M+H]^+^ of ‘metabolite 5’, in positive ionization mode revealed an isotope distribution typical for chlorination. The relative abundance of the +2 and +4 isotopes at a ratio of 9:6:1 indicated the presence of two chlorine atoms for altogether six out of nine metabolites. For metabolites 3, 7 and 14, a ratio of 10:10:3:1 indicated three chlorine atoms. **b**, MS/MS spectrum of *m*/*z* 587.0295, [M-H]^-^ of ‘metabolite 5’, in negative mode revealed iodination by an intense iodide fragment with *m*/*z* 126.90 and a neutral loss of HI. Neutral losses of both HI and HCl are prominent, and were also confirmed by MS/MS analysis in positive mode (Supplementary Data). **c**, Extracted ion chromatograms (EIC) of the iodide fragment in the bag with lowest (bag 3; grey line) and highest (bag 4; black line) viral abundance as present at the end of the *E. huxleyi* demise phase (day 23). Each peak indicates the presence of an iodine-containing metabolite, highlighting a general metabolic shift towards iodination due to viral infection. Numbered peaks are listed in Fig. 3b and Supplementary Table 7. **d**, Peak area profiles reveal the presence of the chloro-iodo metabolites throughout the bloom and demise phase of *E. huxleyi* (separated by dashed line) in bag 3 and bag 4. Peak areas were normalized and scaled to day 23 of bag 4. **e**, Chloro-iodo metabolites were detected in biomass from oceanic *E. huxleyi* blooms in the North Atlantic with higher abundances in ‘late infection’ compared to ‘post infection’ bloom stages. Peak areas were normalized and scaled to cast 29 at 5 m depth.

We further investigated general differences in the occurrence of iodine-containing metabolites as a function of viral infection and revealed additional metabolites in bag 4 (Fig. 4c), indicating a general metabolic shift towards iodination during virus-induced bloom demise. The chloro-iodo metabolites were not detected in the fjord water and in the absence of *E. huxleyi*, further highlighting their close association with viral infection of *E. huxleyi* (Extended Data Fig. 5). Examination of their temporal profiles indicated an early appearance, already one day before the first detection of extracellular EhV, which preceded the onset of bloom demise by two days (day 16 in bag 4; Fig. 4d). The abundance of chloro-iodo metabolites increased further during the demise phase until day 20 and then remained constant until day 23, indicating that these halogenated vDOM metabolites are not immediately consumed, transformed or exported. Thus, these chloro-iodo metabolites may function as sensitive biomarkers for virus-induced demise in the natural environment.

### Chloro-iodo metabolites as hallmarks of viral infection in the ocean

We further sought to examine the significance of the chloro-iodo metabolites as metabolic signatures of viral infection in oceanic *E. huxleyi* blooms. Biomass samples for metabolite analysis of *E. huxleyi* blooms at different infection stages were collected during the NA-VICE cruise in the North Atlantic^29^. The infection stages were characterized using several diagnostic markers: ‘late infection’ sites were defined by high EhV *mcp* levels, whereas at ‘post infection’ sites only low levels of *E. huxleyi* were detected, indicating that samples were collected after bloom demise^29^. We were able to detect high intensities of five chloro-iodo metabolites in the endometabolome of the ‘late infection’ site (cast 29; Fig. 4e). In contrast, only trace amounts of these metabolites were detected in the two ‘post infection’ sites (cast 25 and 27), likely as a result of host lysis, which shunts these metabolites to the DOM pool. The presence of the chloro-iodo metabolites in biomass samples further suggests that their formation is directly associated with infected *E. huxleyi* cells, or alternatively, with microbial organisms that are closely related to this host-virus interaction. This discovery reveals the distinct halogenated metabolic signature of viral infection of *E. huxleyi* blooms in the ocean.

Halogenation is a prominent attribute of natural products in the marine environment due to the high concentrations of halide ions in seawater, with ∼500 mM chloride, ∼1 mM bromide and ∼0.001 mM iodide^30^. Accordingly, several thousand chlorine- and bromine-containing metabolites have been reported, compared to less than 200 iodine-containing metabolites^31^. Among them are only a small number of chloro-iodo metabolites, including tryptophan derivatives isolated from a marine sponge^32^ and coral-derived diterpenoids^33^. In the pelagic ecosystem, the formation of metabolites with a halogenation pattern of both chlorination and iodination is rare. While some halogenated metabolites possess antioxidant and anti-pathogen activity^34^, little is known about their ecological roles in the marine environment. All known algal halogenases belong to the class of haloperoxidases^35^, which lead to the formation of halogenated metabolites following the oxidation of halides by H_2_O_2_. Recently, H_2_O_2_ was shown to play a pivotal role in regulating *E. huxleyi* cell death during the onset of lytic viral infection^36^. Therefore, the increased levels of halogenated metabolites may be a result of enhanced haloperoxidase activity in *E. huxleyi* cells, acting as a possible strategy to scavenge surplus ROS during viral infection. Recently, a halogenase encoded by a marine cyanophage was discovered^37^, illustrating how viral infection can expand the metabolic capabilities of the infected host. Following cell lysis, the chloro-iodo metabolites may serve as a carbon source for specialized microbial taxa that encode dehalogenases, such as the phytoplankton-associated *Rhodobacteraceae*^45^.

The importance of the ‘viral shunt’ to the marine DOM pool was first highlighted more than 20 years ago^4^. Since then, the ubiquitous presence of viruses in the ocean has become increasingly apparent^38^. Nevertheless, we still lack quantitative tools to assess the extent of the ‘viral shunt’, its metabolic composition, and the direct consequences for the marine microbial community. By applying an untargeted exometabolomics approach in high temporal resolution, we mapped the metabolic landscape of biologically active, labile DOM that evolves during phytoplankton bloom succession. We further resolved the metabolic footprint of lytic viral infection in the marine environment, revealing halogenation as a hallmark of the *E. huxleyi* vDOM. The semi-stable nature of the halogenated vDOM metabolites may enable their application as metabolic biomarkers for sensitive quantification of the extent of the ‘viral shunt’ in phytoplankton blooms. Importantly, the increased stability of halogenated vDOM in the marine environment illustrates how lytic viral infection can lead to rapid carbon sequestration in the ocean via the microbial carbon pump^39^. Decoding the unique metabolic landscape produced by different microbial interactions that control cell fate in phytoplankton blooms, such as host-pathogen interactions, will provide essential insights into the impact of these microscale interactions on large-scale biogeochemical processes in the marine environment.

## Methods

### Chemicals and internal standards

All solvents and metabolite standards were obtained at highest purity. Methanol (Chromasolv LC-MS Ultra) used for solid phase extraction and hydrochloric acid (HCl, ≥32% (T), Fluka) were purchased from Honeywell (Seelze, Germany). For all other purposes, methanol (Ultra gradient HPLC) was purchased from J. T. Baker (Norway). Acetonitrile (ULC/MS), methyl tert-butyl ether (MTBE, HPLC), hexane (HPLC) and formic acid (>99%, ULC/MS) were purchased from Bio-Lab (Jerusalem, Israel). Acetone (≥ 99.8%, Chromasov HPLC) was purchased from Sigma-Aldrich (Saint Louis, MO, USA). Water (HiPerSolv Chromanorm) used for solid phase extraction was purchased from VWR (Oslo, Norway). For all other purposes, water was purified by a Milli-Q system (resistivity 18.2 MΩ cm at 25°C, TOC < 5 ppb, Merck Millipore, Molsheim, France). Indole-3-acetic-2,2-d_2_ acid (≥ 98%) and *N*-hexanoyl-L-homoserine lactone-d_3_ (≥ 99%) were used as isotopically labelled extraction standards (both from Santa Cruz Biotechnology, Dallas, TX, USA). Caffeine-(trimethyl-d_9_) (98%, Sigma-Aldrich) and L-tryptophan-d_5_ (98%, Cambridge Isotope Laboratories, Andover, MA, USA) were used as isotopically labelled injection standards for ultra performance liquid chromatography-high-resolution mass spectrometry (UPLC-HRMS) analysis.

### Mesocosm setup and sampling

The mesocosm experiment AQUACOSM VIMS-Ehux was carried out between 24^th^ May (day 0) and 16^th^ June (day 23) 2018 in Raunefjorden at the Marine Biological Station Espegrend, Norway (60°16′11N; 5°13′07E) as previously described^40^. Four light-transparent enclosure bags were filled with nonfiltered surrounding fjord water (day -1; pumped from 5 m depth) and continuously mixed by aeration (from day 0 onwards). Each bag was supplemented with nutrients at a nitrogen to phosphorous ratio of 16:1 (1.6 µM NaNO_3_ and 0.1 µM KH_2_PO_4_ final concentration) on days 0-5 and 14-17, whereas on days 6, 7 and 13 only nitrogen was added. Nutrient concentrations were measured daily^41^. Water samples were collected daily (07:00 AM) from each bag and the surrounding fjord, which served as an environmental reference. For flow cytometry, water samples were collected in 50 mL tubes from approximately 1 m depth. For all other purposes, water samples were collected in 10-20 L carboys (rinsed with <100 kDa filtered seawater) from approximately 1 m depth using a peristaltic pump at ca. 5 L/min and pre-filtered with a 200 µm nylon mesh. Samples were kept at 4°C until further processing.

### Enumeration of phytoplankton cells by flow cytometry

Water samples were pre-filtered using 40 µm cell strainers and immediately analysed with an Eclipse iCyt flow cytometer (Sony Biotechology, Champaign, IL, USA) as previously described^21^. A total volume of 300 µL with a flow rate of 150 µL/min was analyzed. A threshold was applied on the forward scatter to reduce background noise. Four phytoplankton populations were identified by plotting the autofluorescence of chlorophyll (em: 663-737 nm) versus phycoerythrin (em: 570-620 nm) and side scatter: calcified *E. huxleyi* (high side scatter), *Synechococcus* (high phycoerythrin), nano-(high chlorophyll) and picophytoplankton (low chlorophyll; Extended Data Fig. 2).

### Enumeration of extracellular EhV by qPCR

Water samples (1-2 L) were sequentially filtered by vacuum through hydrophilic polycarbonate filters with a pore size of first 20 µm (47 mm; Sterlitech, Kent, WA, US), then 2 µm (Isopore, 47 mm; Merck Millipore, Cork, Ireland), and finally 0.22 µm (Isopore, 47 mm; Merck Millipore). Filters were immediately flash-frozen in liquid nitrogen and stored at -80°C until further processing. DNA was extracted from the 0.22 µm filters using the DNeasy PowerWater kit (QIAGEN, Hilden, Germany) according to the manufacturer’s instructions. Each sample was diluted 100 times, and 1 µL was then used for qPCR analysis. EhV abundance was determined by qPCR for the major capsid protein (*mcp*) gene^42^: 5′-acgcaccctcaatgtatggaagg-3′ (mcp1F^43^) and 5′-rtscrgccaactcagcagtcgt-3′ (mcp94Rv; Mayers, K. et al., unpublished). All reactions were carried out in technical triplicates. For all reactions, Platinum SYBER Green qPCR SuperMix-UDG with ROX (Invitrogen, Carlsbad, CA, USA) was used as described by the manufacturer. Reactions were performed on a QuantStudio 5 Real-Time PCR System equipped with the QuantStudio Design and Analysis Software version 1.5.1 (Applied Biosystems, Foster City, CA, USA) as follows: 50°C for 2 min, 95°C for 5 min, 40 cycles of 95°C for 15 s, and 60° C for 30 s. Results were calibrated against serial dilutions of EhV201 DNA at known concentrations, enabling exact enumeration of viruses. Samples showing multiple peaks in melting curve analysis or peaks that were not corresponding to the standard curves were omitted.

### Sampling and extraction of dissolved organic matter by solid phase extraction

To collect dissolved organic matter (DOM) of <0.22 µm particle size, water samples were filtered gently, acidified, and extracted by solid phase extraction (SPE). Glassware and chemically resistant equipment were used whenever possible and cleaned with HCl (1% or 10%) and Deconex 20 NS-x (Borer Chemie, Zuchwil, Switzerland) to reduce contaminations. On day 5, no samples were extracted for DOM analysis. Water samples were first gravity-filtered over 25 µm stainless steel filters (47 mm, Sinun Tech, Barkan, Israel), which were pre-cleaned by thorough washing in a polarity gradient of organic solvents (water, methanol, acetone, hexane). Filtrates were then filtered gently by vacuum (<400 mbar under-pressure) over pre-combusted GF/A filters (≥ 5 h at 460°C; 47 mm, GE Healthcare Whatman, Buckinghamshire, UK), and finally over 0.22 µm pre-cleaned^44^ hydrophilic PVDF filters (<600 mbar under-pressure; 47 mm, Durapore, Merck Millipore, Cork, Ireland). Filtration led to 70-84% reduction in the abundance of large virus-like particles (VLP) and to the complete removal of bacteria (Extended Data Fig. 6). Per sample, 1 L of filtrate was collected in a glass bottle and supplemented with 5 µL internal standard solutions containing *N*-hexanoyl-L-homoserine lactone-d_3_ (0.2 µg/µL in methanol) and indole-3-acetic-2,2-d_2_ acid (1 µg/µL in water), except for days 1-2, in which no internal standards were added. Filtrates were incubated for 2-3 hours at 4°C in the dark and then acidified to pH 2 using 10% HCl^45^. Metabolites were extracted using hydrophilic-lipophilic balanced SPE cartridges (Oasis HLB, 500 mg, Waters, Milford, MA, USA) as follows: cartridges were conditioned (6 mL methanol), equilibrated (6 mL 0.01 N HCl), and then loaded by gravity with the acidified samples (1.5-2.5 h). The cartridges were then washed (18 mL 0.01 N HCl), dried completely using a vacuum pump, and gravity-eluted with 5 mL methanol into 4 mL glass vials. In total, 110 biological samples were collected. Blank samples and internal standard quality control (IS QC) samples were obtained every 5 days (in total 19 samples; Supplementary Table 1). Eluates were kept at -80°C, dried under a flow of nitrogen (TurboVap LV, Biotage, Uppsala, Sweden) within 1.5 months after collection, and stored at -80°C until further processing.

### Untargeted profiling of semi-polar metabolites by UPLC-HRMS

Biological samples were randomized and divided into three batches with approximately 40 samples in each batch, including blanks and IS QC samples (Supplementary Table 2). Randomization was performed automatically using an in-house R^46^ script with the following constraints: the total number of biological samples per batch was either 36 or 37, of which 7 or 8 samples were randomly sampled from the pool of fjord samples and the remaining samples were randomly sampled from the pool of bag samples. Every experimental sampling day was represented at least once in each analytical batch. Per batch, samples were thawed, re-dissolved in 310 μL methanol:water (1:1, ν:ν) containing tryptophan-d_5_ (2.1 µg/mL) and caffeine-d_9_ (1.5 µg/mL) as injection standards, vortexed, sonicated for 10 min, and centrifuged at 3,200×g for 10 min at 4°C. The supernatants were transferred to 200 μL glass inserts in autosampler vials and directly used for LC-MS analysis. An aliquot of 1 µL was analyzed using UPLC coupled to a photodiode detector (ACQUITY UPLC I-Class, Waters) and a quadrupole time-of-flight (QToF) mass spectrometer (SYNAPT G2 HDMS, Waters), as described previously^47^ with slight modifications. Briefly, metabolites were separated using an ACQUITY UPLC BEH C18 column (100 × 2.1 mm, 1.7 µm; Waters) attached to a VanGuard pre-column (5 × 2.1 mm, 1.7 µm; Waters) with a gradient of 5-100% acetonitrile at a flow of 0.3 mL/min and a total run time of 40 min. The mobile phases consisted of 0.1% formic acid in either acetonitrile:water (5:95, ν:ν, mobile phase A) or acetonitrile (mobile phase B). The chromatographic gradient was set to linear from 100% to 72% mobile phase A over 22 min and from 72% to 30% mobile phase A over 13.5 min, after which the column was first washed with 100% mobile phase B for 2 min and then returned to initial conditions (100% mobile phase A) and equilibrated for 1.5 min. The PDA detector was set to 200-600 nm. A divert valve (Rheodine) excluded 0-1.2 min and 35-40 min from injection to the mass spectrometer. The ESI source was set to 140°C source and 450°C desolvation temperature, 1.5 kV capillary voltage, and 20 eV or 27 eV cone voltage (positive or negative ionization mode, respectively), using nitrogen as desolvation gas (800 L/h) and cone gas (53 L/h). The mass spectrometer was operated in full scan MS^E^ positive and negative resolution mode (26,000 at *m*/*z* 556) over a mass range of 50-1600 Da alternating with 0.1 min scan time between low-(1 eV collision energy) and high-energy scan function (collision energy ramp of 10-45 eV in positive and 10-40 eV in negative ionization mode). LC-MS analyses were performed over two weeks with about 120 injections per batch in the positive and negative ionization mode including blanks, IS QC, authentic standard mixtures, reference material, and aliquots of a pooled QC sample (Supplementary Tables 3 and 4). The pooled QC sample was generated by combining aliquots of 10 µL from all biological samples of the first batch. For each batch, a new aliquot was transferred into an injection vial.

### Comparative analysis of untargeted metabolite profiling data

LC-MS files were converted from Waters RAW binary files to open-format ‘mzXML’ files using the command line version of the ‘msconvert’ utility as part of the ProteoWizard toolkit^48^. Conversion parameters were set as follows: 64-bit numeric accuracy, ‘zlib’ compression enabled, and ‘scanEvent’ filter set to ‘1’ corresponding with the full-scan acquisition channel. Pre-processing of the ‘mzXML’ files, which generates matrices with aligned LC-MS features across experimental samples with corresponding integrated peak area values, was done in R^46^ using the packages ‘xcms’^49^ and ‘CAMERA’^50^ obtained from the Bioconductor repository (www.bioconductor.org). More recent 3.x versions of ‘xcms’ have dedicated functions for quality control of raw data pre-processing, for which parameters were fine-tuned (Supplementary Tables 5 and 6; Extended Data Fig. 7). Feature matrices were inter-and intra-batch corrected using an in-house R script applying the algorithm presented by van der Kloet *et al*.^51^. Briefly, peak intensities in each batch were corrected for systematic variations in MS sensitivity by using a non-linear curve fitted to peaks from the pooled QC samples, whereas non-systematic inter-batch fluctuations in MS sensitivities were corrected by adjusting the medians of the regression curves. Principal components analysis (PCA) was applied to the batch-corrected data to have an initial overview of the sample separation and as a reference for the data normalization procedure. Two fjord samples (day 10 and day 19) were identified as outliers and omitted from further analysis. Choice of the most appropriate normalization method was based on the following experimental constraints: absence of technical replicates, potentially high variability between biological groups (the mesocosm bag samples are not true replicates), and the strong influence of background environmental fluctuations. The probabilistic quantile normalization (PQN) algorithm^52^ was chosen as it applies minimal assumptions about the data and instead relies on the empirical distribution in a reference sample set. Each peak intensity is thereby normalized by the quotient of the same peak in the reference sample (the average of the pooled QC samples). As pooled QC samples capture most of the analytical variation that remains after batch correction and some of the environmental variability, they are a natural choice as reference sample set. PCA was then re-applied to the normalized feature matrix in the positive (Fig. 1c) and the negative (Extended Data Fig. 8) ionization modes.

One-way repeated-measures ANOVA was performed to test if the DOM composition, as represented by the scores of PC axis 1, changed significantly through time. Using the R package ‘nlme’, a linear mixed-effects model was fitted by setting the factor time as fixed effect and the factor bag number as random effect. The degree of change in DOM composition was assessed for each phytoplankton bloom phase by fitting a liner regression model using the R function ‘lm’. Furthermore, a Pearson correlation analysis was performed to correlate PCA axis 1 with the parameters water temperature and flow cytometry-based chlorophyll levels using the R function ‘cor’, while omitting missing values.

Hierarchical cluster analysis was applied to the feature matrix in the positive ionization mode. First, data was filtered by removing features in which the intensity of the fjord samples was higher than 60% of the maximum intensity for that particular feature. While being a very simplistic filtering approach, it removed most features related to environmental changes and allowed us to focus on features that are most relevant to the biological processes of interest. The feature matrix was then log-transformed and standardized per mass feature. The ‘heatmap.2’ function from the ‘gplots’ R package was used to generate a heatmap with row-wise scaling and clustering and with the ‘redblue’ colour panel. Columns were ordered according to the experimental day factor to preserve the temporal property of the data and observe possible trends over time.

### Correlation analysis of metabolite profiling data with extracellular EhV abundance

To focus on the bloom and demise phases of *E. huxleyi* as indicated by cell enumeration data, the raw data pre-processing steps were re-applied as described above, however, for a reduced number of samples (days 10 to 23) with more sensitive peak grouping parameters (‘minSamples=8’, ‘minFraction=0’; Supplementary Tables 5 and 6). This re-iteration enabled the detection of features that were present in some of the mesocosm bags and for a few days only. The feature matrices resulting from this step contained almost twice as many features as the first, global feature matrices, and were used, together with the extracellular EhV abundance data, for correlation and differential analysis. Pearson’s correlation coefficient values between the temporal profile of extracellular EhV and each feature corresponding with mesocosm bag 4 were calculated using the ‘cor.test’ function in the R programming language. The ‘alternative’ was set to ‘greater’ to find only features that are positively correlated with extracellular EhV abundance. Features were filtered according to the following thresholds: estimated correlation >0.7; *P* <0.05, and intensity of feature in the fjord samples <60% of the maximum intensity in the mesocosm bag samples, resulting in 705 features. The log-transformed peak intensities were clustered using the ‘agnes’ function in the R package ‘cluster’ and plotted as a heatmap using the function ‘heatmap.2’ (Extended Data Fig. 4). Columns were ordered according to each mesocosm bag to highlight the differences between them. A sub-cluster of 141 features showed distinct differences between bag 3 and bag 4. Individual intensity profiles of each feature were plotted for the mesocosm bags and the fjord across days 10 to 23. These plots were used for manual inspection, which allowed the selection of a subset of 65 features that showed differential intensities between mesocosm bags as observed for EhV abundance. Manual curation of these features revealed 20 feature groups in which the isotopes, adducts and in-source fragments were annotated (Supplementary Data). Finally, a heatmap of the most intense feature of each group (the molecular ion adduct or the water loss fragment) was plotted using the function ‘heatmap.2’ and the ‘PuBuGn’ colour panel (‘RColorBrewer’ R package). The columns were ordered first according to days and then bags following their level of viral infection, and the rows according to retention time.

### Structural characterization of the vDOM metabolites

For structural information of the 20 feature groups, MS/MS analyses were performed for the putative molecular ion of each feature group both in positive and in negative ionization mode using a collision energy ramp of 10-45 eV and scan time of 0.5 sec. Analyses were performed on samples with high intensities, namely samples from day 22 and day 23 of mesocosm bag 4. In case of low signal intensity of the molecular ion, up to 3 µL were injected, or the most intense fragment ion ([M+H-H_2_O]^+^) or adduct ion ([M-H+FA]^-^) was selected, if available. Manually curated MS/MS spectra were used to annotate fragments and neutral losses and for the prediction of molecular formulas using SIRIUS 4.0.1^53^. For formula prediction, the following elements were allowed: 0-3 chlorine atoms (based on isotopic distribution), 0-1 iodine atom (based on presence of *m*/*z* 126.90 in MS/MS spectra acquired in negative ionization mode), 0-infinite nitrogen atoms (set to ‘0’ if only odd fragments were detected). Default settings were used for C, P, H, S and O (‘0-infinite’). Mass accuracy was set to 10 ppm. For annotated mass spectral information, see Supplementary Data file. The predicted molecular formulas of the chlorine-iodine containing metabolites were searched against several external mass spectral and natural product databases, including DEREP-NP^54^ and COCONUT^55^ (for a full list see Supplementary Table 7). The occurrence of iodine-containing metabolites in bags 3 and 4 at the end of the *E. huxleyi* demise phase (day 23) was compared by plotting the extracted ion chromatogram (EIC) of the iodide fragment (*m/z* 126.90) in negative ionization mode using the high-energy scan function (MassLynx, version 4.1, Waters).

### Extraction and metabolite profiling of *E. huxleyi* blooms in the North Atlantic

Water samples of natural *E. huxleyi* blooms were collected during the NA-VICE cruise as described previously^29^. To extract metabolites that are associated with viral infection, cast 29 was selected, which represents a ‘late infection’ stage^56^, in addition to casts 25 and 27, which represent ‘post infection’ stages^29,56^. Samples (1-1.5 L) were pre-filtered through 200 µm mesh and collected on 0.8 µm hydrophilic polycarbonate filters (47 mm, Millipore). The filters were then flash-frozen in liquid nitrogen and stored at -80°C until processing seven years post collection. Extraction was performed as described previously^21^ with slight modifications. Briefly, filters were extracted with 3 mL pre-cooled (−20°C) methanol:MTBE (1:3, ν:ν) solution containing *N*-hexanoyl-L-homoserine lactone-d_3_ (0.33 µg/mL) and indole-3-acetic-2,2-d_2_ acid (1.67 µg/mL) as internal standards. The samples were shaken for 30 min at 4°C and sonicated for 30 min. The samples were then supplemented with 1.5 mL water:methanol (3:1, ν:ν) solution, vortexed for 1 min, and centrifuged for 10 min at 3,200×g and 4°C. The upper organic phase was removed (1.5 ml) and the remaining aqueous phase was re-extracted with 1.5 mL pre-cooled MTBE to further reduce lipophilic metabolites. Finally, the aqueous phase (1.5 mL) was transferred into 2 mL centrifuge tubes, centrifuged at 28,000×g, and the supernatant was dried under a flow of nitrogen (TurboVap LV) followed by lyophilization (Gamma 2-16 LSCplus, Martin Christ, Osterode am Harz, Germany). The dried extracts were kept at -80°C until analysis. Untargeted profiling of semi-polar metabolites by UPLC-HRMS analysis was performed as described above, using an aliquot of 3 µL of each re-suspended extract.

### Temporal profiles of chloro-iodo metabolites

Temporal intensity profiles of the chloro-iodo metabolites were generated using MassLynx and QuanLynx (Version 4.1, Waters). Peak areas were extracted for the [M+H]^+^ or [M+H-H2O]^+^ ions above a signal-to-noise threshold of 10. The peak areas were then normalized to the extraction standard indole-3-acetic-2,2-d_2_-acid. Samples from the NA-VICE cruise were further normalized to the filtered volume. Analysis of blank samples from the mesocosm experiment (n = 10; Supplementary Table 1) ruled out that any of the metabolites originated from sample processing.

## Data availability

Data supporting the findings of this study are available in the paper and its Supplementary Information files. Flow cytometry data and nutrient data are available in Dryad^41^.

## Acknowledgments

We thank all team members of the VIMS-Ehux project for setting up and conducting the mescosom experiment, especially Jorun Egge, Aud Larsen, Tatiana Tsagaraki, Celia Marrase and Rafel Simó. We are grateful to Ilana Rogachev for technical support on the LC-MS instrument. We further thank the team members and crew of the NA-VICE cruise for assistance at sea, as well as the Marine Facilities and Operations at the Woods Hole Oceanographic Institution for logistical support. Finally, we acknowledge all comments on the manuscript draft received from Roy A. Meoded and Tim U. H. Baumeister. This research was supported by the European Research Council CoG (VIROCELLSPHERE grant no. 681715) and a research grant from the Estate of Bernard Berkowitz awarded to A.V., and the Dean of Faculty Fellowship of the Weizmann Institute of Science awarded to C.K. The mesocosm experiment VIMS-Ehux was supported by EU Horizon2020-INFRAIA project AQUACOSM (grant no. 731065). The NA-VICE cruise was supported by the National Science Foundation (grant no. OCE-1059884).

## Author contributions

C.K., G.S. and A.V. conceptualized the project and wrote the manuscript. C.K. and G.S. designed and performed the experiments. N.S., C.K. and G.S. performed the LC-MS data analysis. F.V. acquired the flow cytometry data. D.S. and G.S. extracted DNA and performed the qPCR analysis. All authors commented on the manuscript.

## Additional information

Supplementary Information is available for this paper.

## Extended Data

**Extended Data Fig. 1:**
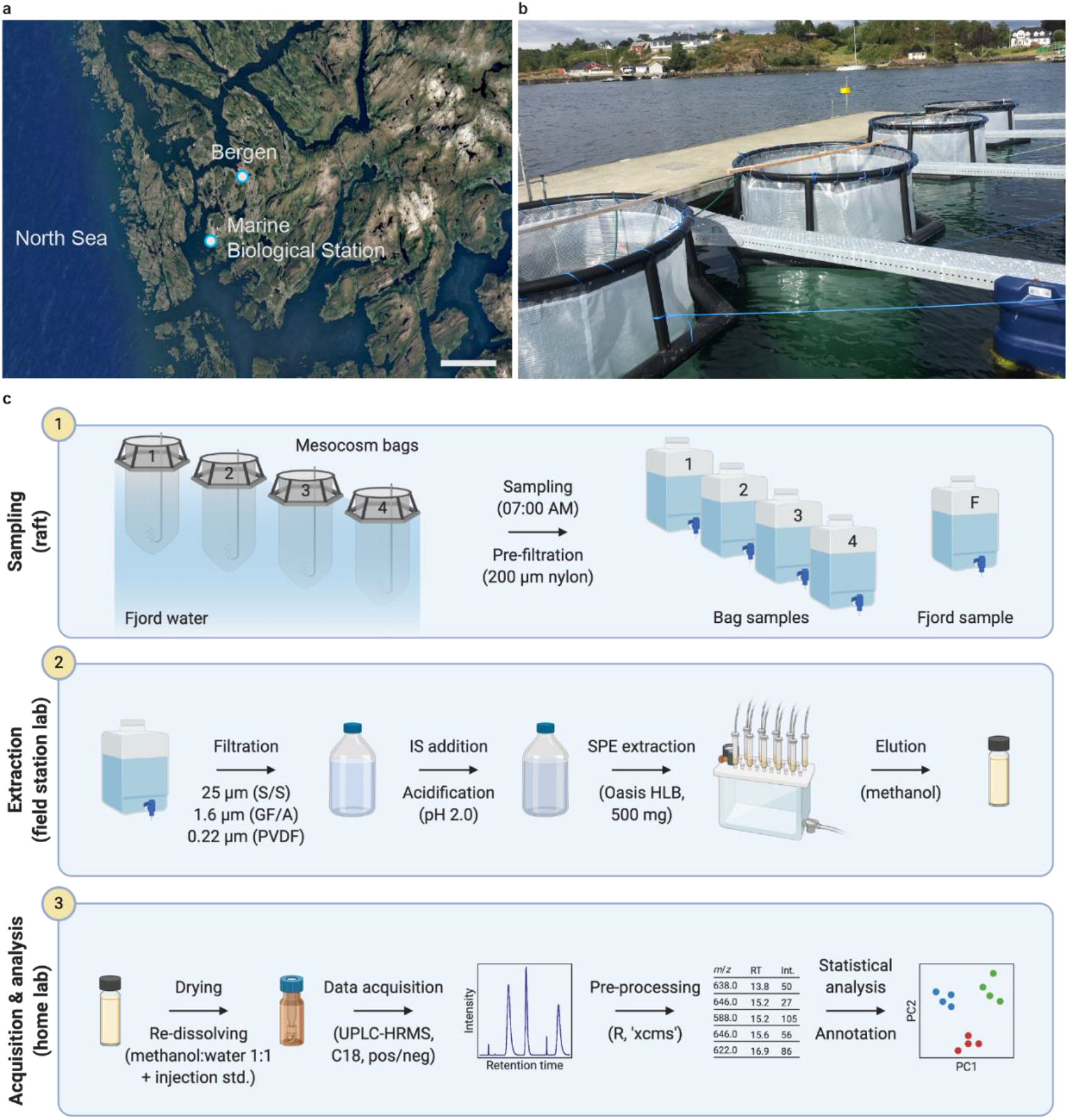
Exometabolomics workflow applied on induced algal blooms in mesocosm setups. **a**, Location of the Marine Biological Station Espegrend, Norway (60°16′11N; 5°13′07E). Scale bar: 10 km. Map data: Google Earth, Landsat/Copernicus. **b**, Experimental setup, consisting of four transparent enclosure bags mounted on floating frames and moored to a raft in the middle of the fjord. The bags were filled with surrounding fjord water at day -1 of the experiment and were supplemented with nutrients throughout the experiment. **c**, Workflow for untargeted exometabolite profiling. Sampling (1): water samples were collected daily (07:00 AM) from each bag and the surrounding fjord using a peristaltic pump and a 200 µm nylon pre-filter. Extraction (2): the water samples were filtered through three subsequent filters (25 µm, 1.6 µm and 0.22 µm). Per sample, 1 L of filtrate was collected in a glass bottle and supplemented with internal standards (IS) for extraction. After 2-3 hours, the filtrate was acidified to pH 2.0. Dissolved metabolites were then extracted with SPE cartridges (Oasis HLB) and eluted with methanol. Eluates were kept at -80°C until further processing. Acquisition and analysis (3): eluates were dried and re-dissolved for untargeted data acquisition using a UPLC-HRMS system. The collected data was pre-processed and the resulting dataset was used for comparative statistical analysis and peak annotation. Workflow created with BioRender.com.

**Extended Data Fig. 2:**
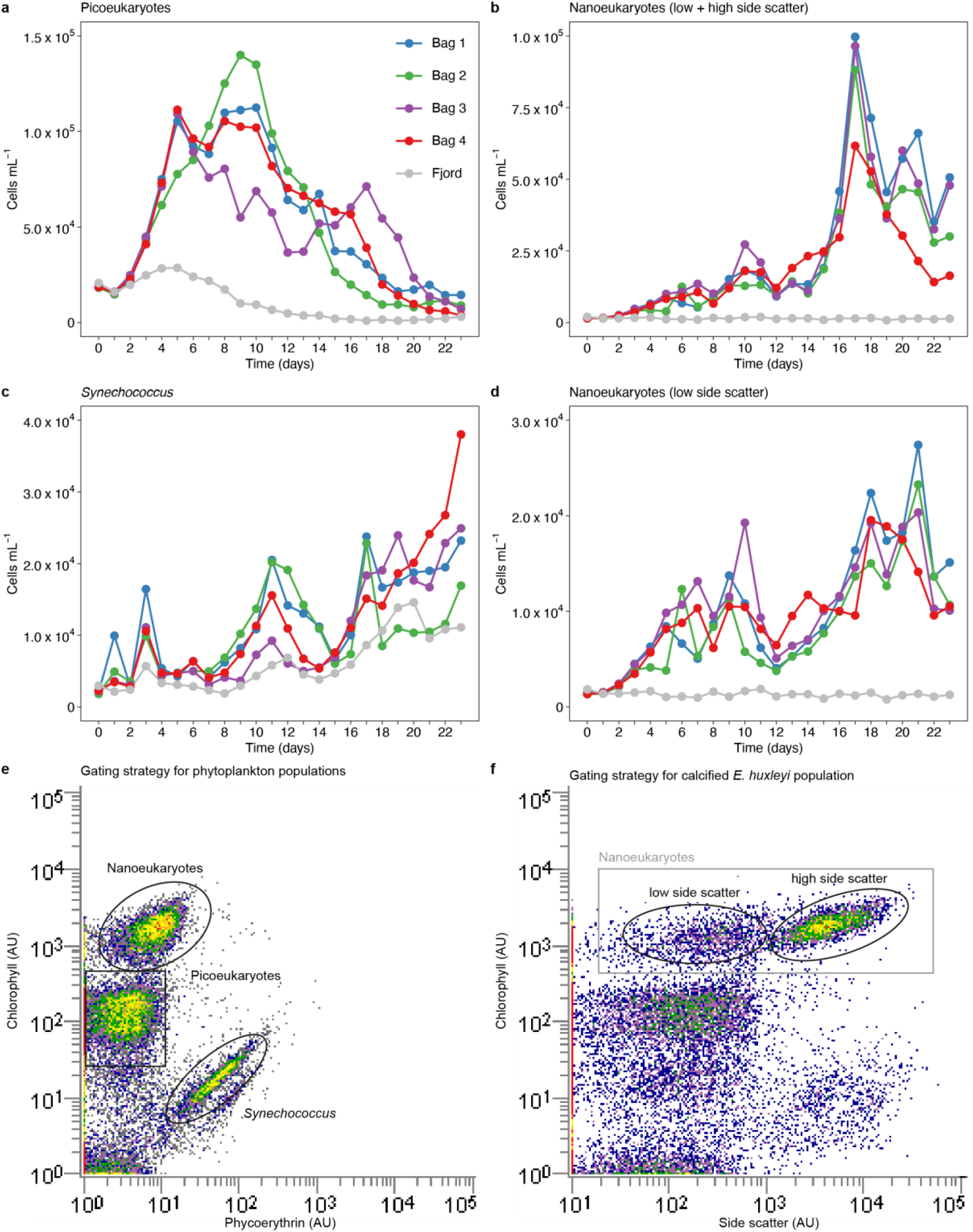
Population dynamics throughout phytoplankton bloom succession in mesocosm setups. Abundance of three phytoplankton populations: picoeukaryotes (**a**), nanoeukaryotes (**b**) and *Synechococcus* (**c**) based on flow cytometry analysis. The populations were identified by plotting the autofluorescence of chlorophyll (em: 663-737 nm) versus phycoerythrin (em: 570-620 nm) (**e**). The nanoeukaryote population was further separated into low side scatter cells (**d**) and high side scatter cells (calcified *E. huxleyi*, as presented in Fig. 1b), by plotting the autofluorescence of chlorophyll versus side scatter (**f**).

**Extended Data Fig. 3:**
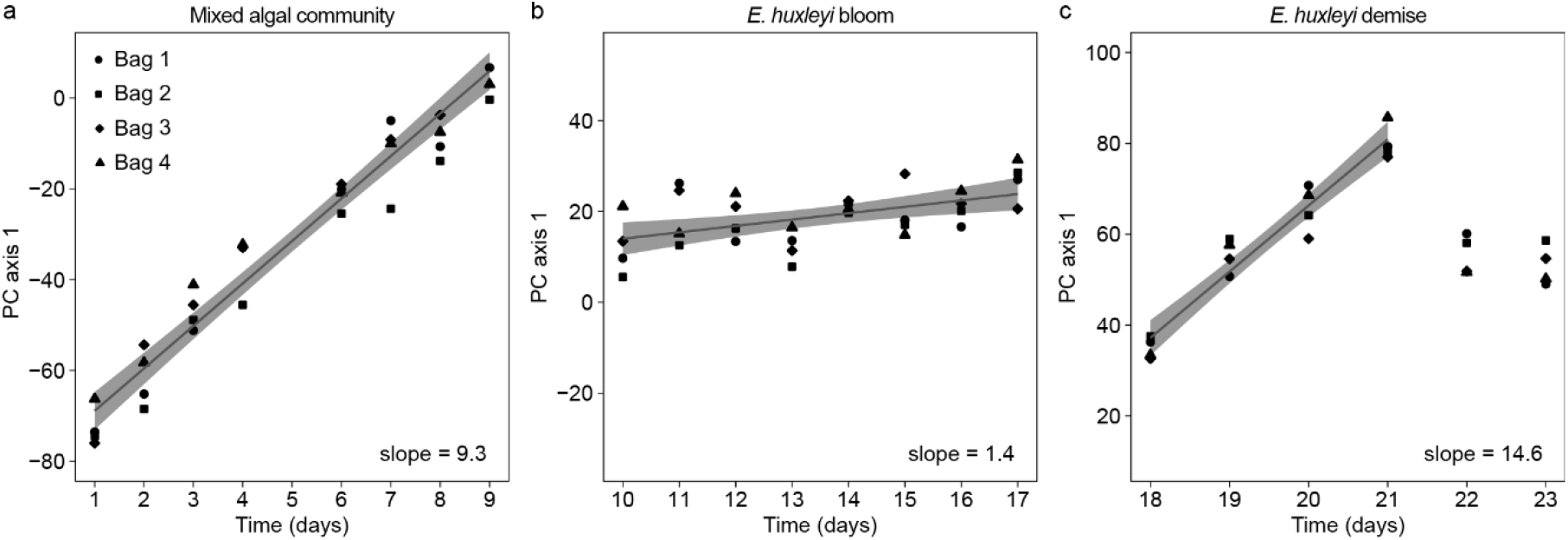
Changes in DOM composition during different phytoplankton bloom phases. The degree of change in DOM composition was assessed for each phase based on the scores of PC axis 1. The initial mixed algal community (**a**) and the *E. huxleyi* demise (**c**) phases presented a high degree of change (slope of 9.3 and 14.6, respectively), while the degree of change during the *E. huxleyi* bloom phase (**b**) was low (slope of 1.4). Days 22-23 were omitted from the linear regression analysis of the demise phase due to a change in directionality. For all linear regression slope estimates, *p* < 0.002.

**Extended Data Fig. 4:**
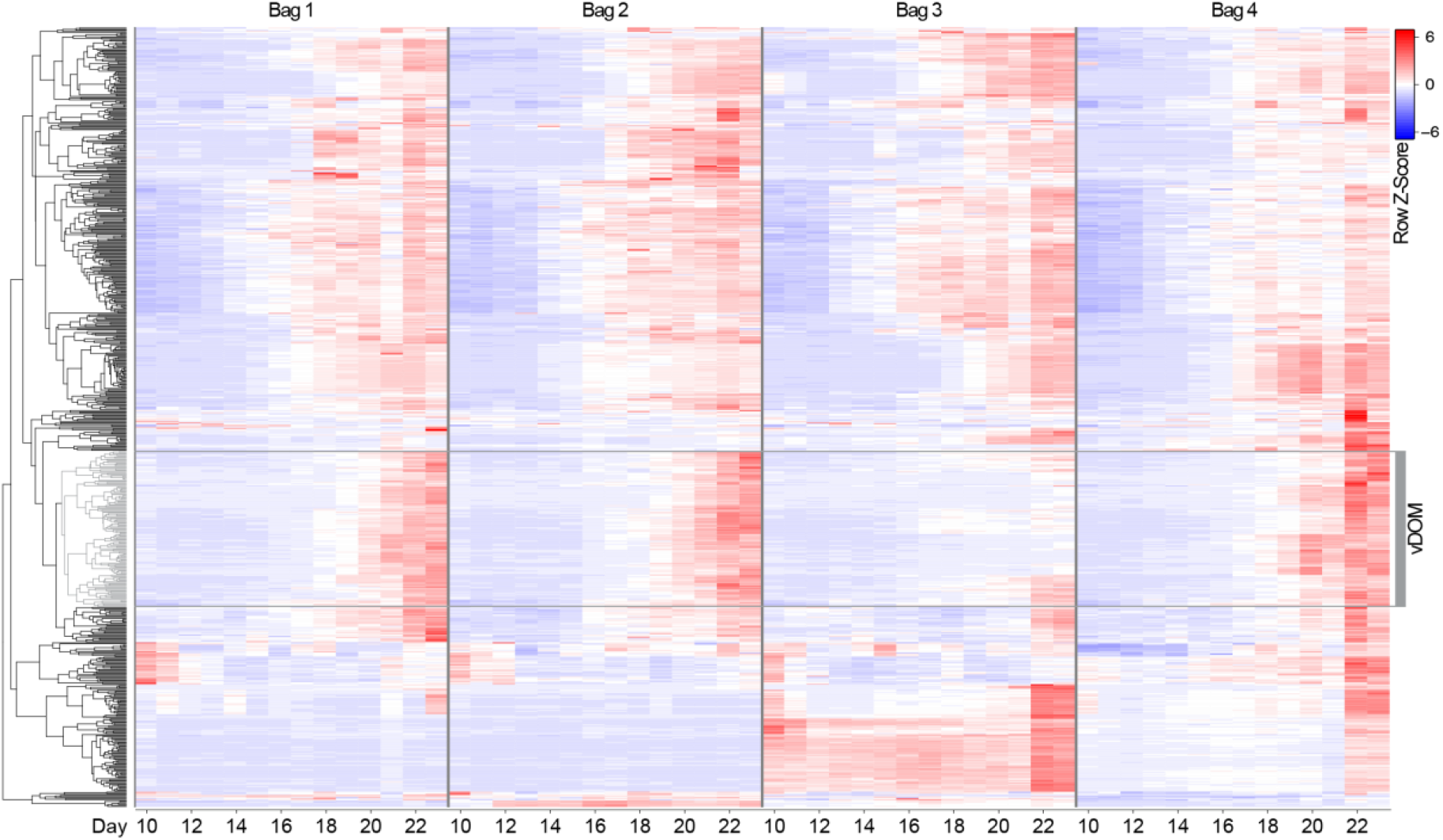
Metabolite features positively correlated with the abundance of extracellular EhV in mesocosm bag 4. Hierarchical cluster analysis of 705 metabolite features with positive correlation (Pearson correlation, *r* >0.7) to the abundance of extracellular EhV during bloom and demise of *E. huxleyi* in bag 4. Samples (columns) are organized per bag, from day 10 to 23. The cluster highlighted in grey contains 141 metabolite features that showed differential abundance between the mesocosm bags, corresponding to the varying degree of viral infection. This cluster, referred to as ‘vDOM’, was selected for further investigation (Fig. 3b).

**Extended Data Fig. 5:**
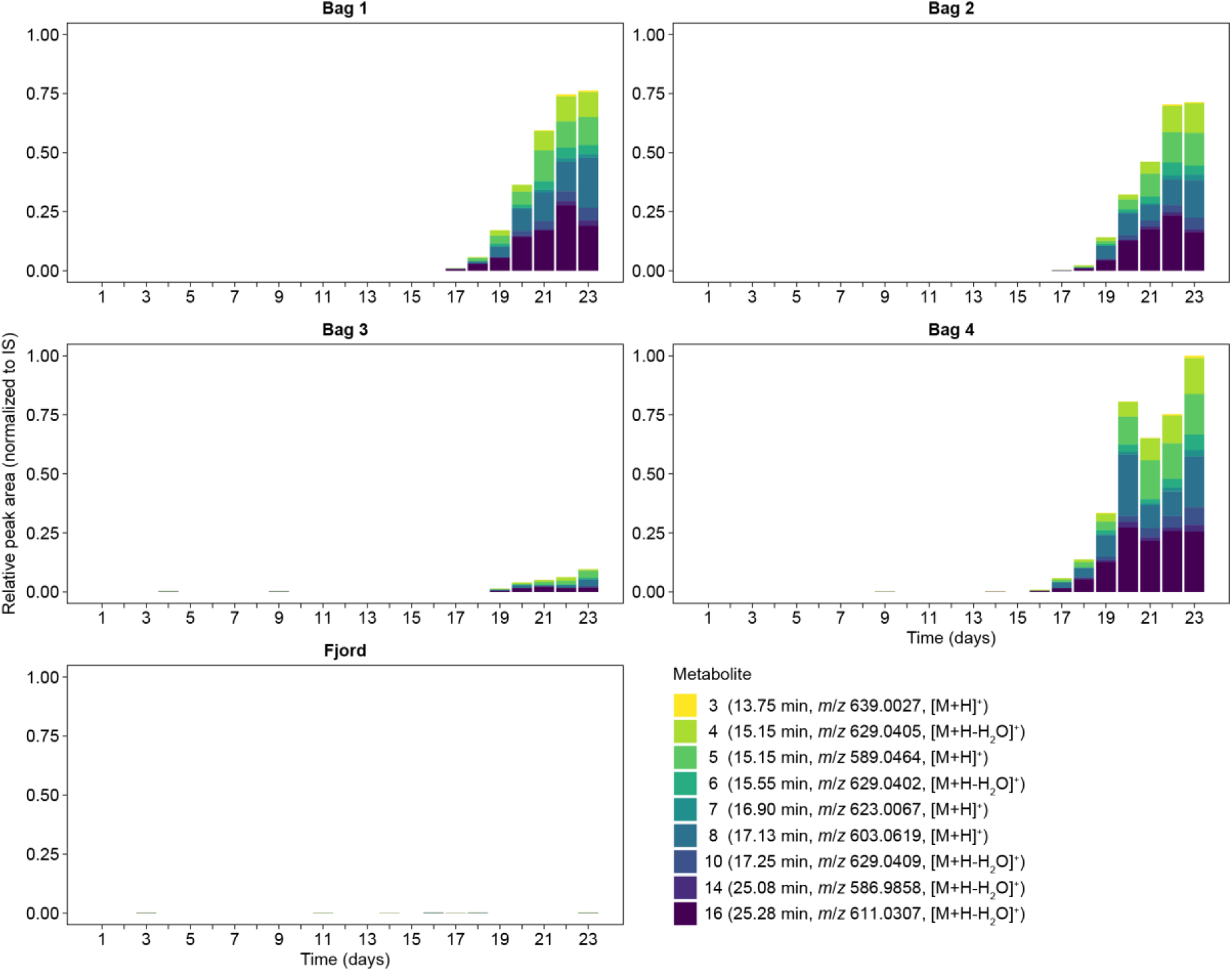
Temporal profiles of nine chlorine-iodine-containing exometabolites in the mesocosm bags and surrounding fjord. Peak areas were integrated for the [M+H]^+^ or [M+H-H_2_O]^+^ ions above a signal-to-noise threshold of 10. The peak areas were then normalized to the extraction standard indole-3-acetic-2,2-d_2_ acid. Graphs are scaled to mesocosm bag 4 at day 23.

**Extended Data Fig. 6:**
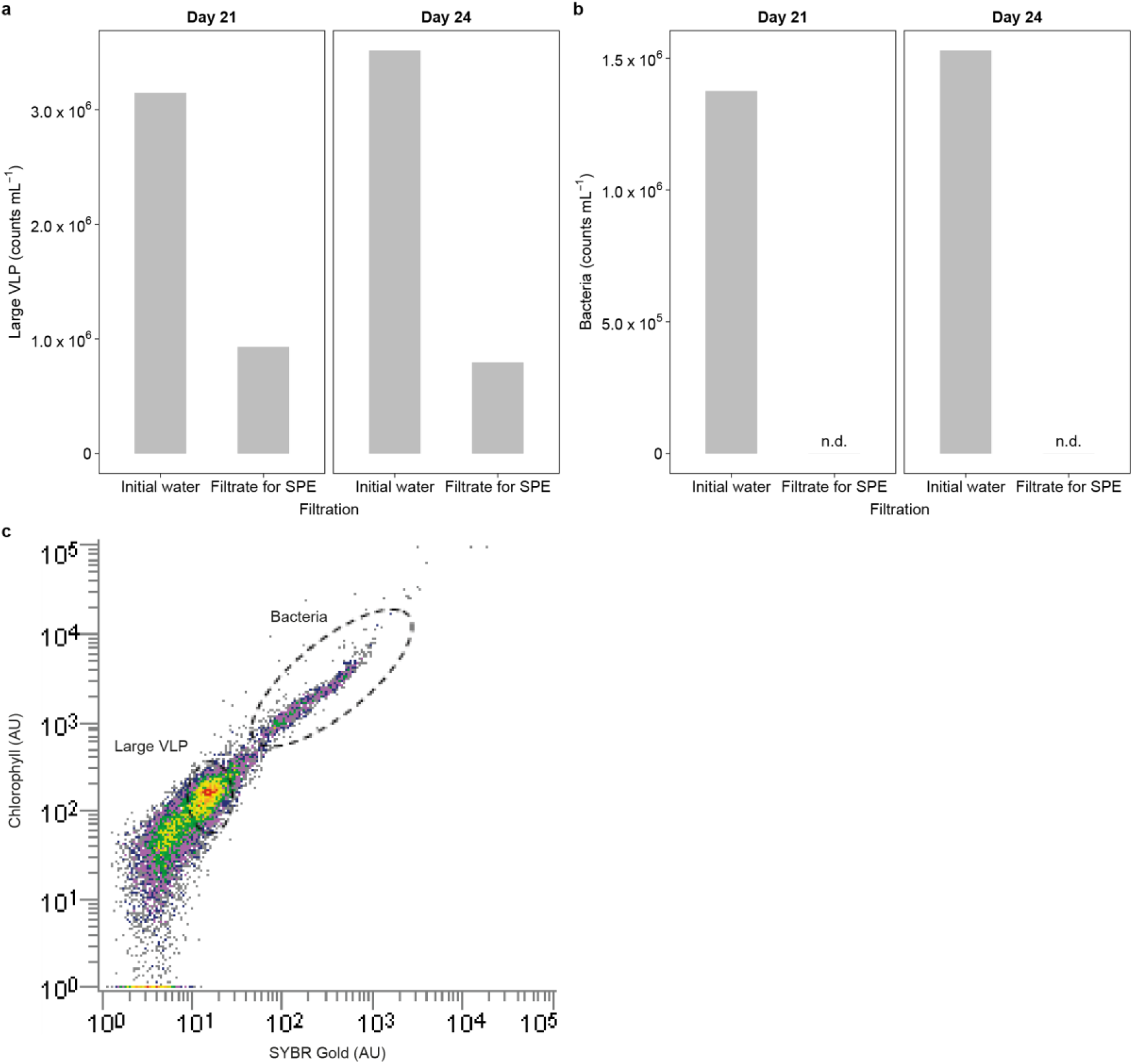
Removal of large virus-like particles (VLP) and bacteria during filtration for exometabolomics. Comparison of the abundance of large VLP (**a**) and bacteria (**b**) in the initial water (after filtration with 25 µm stainless steel filter) and in the filtrate used for SPE (after filtration with 0.22 µm PVDF filter). The comparison was made for samples from bag 4 at two time points during bloom demise (days 21 and 24). Filtration using 0.22 µm filters led to 70-85% reduction in large VLP and to the complete removal of bacteria. Large VLP and bacteria were enumerated by flow cytometry using SYBR Gold staining (**c**, ex: 488 nm, em: 525 nm). The gates were set by comparing to reference samples containing fixed EhV201 and bacteria from lab cultures (see Supplementary Methods). n.d., not detected.

**Extended Data Fig. 7:**
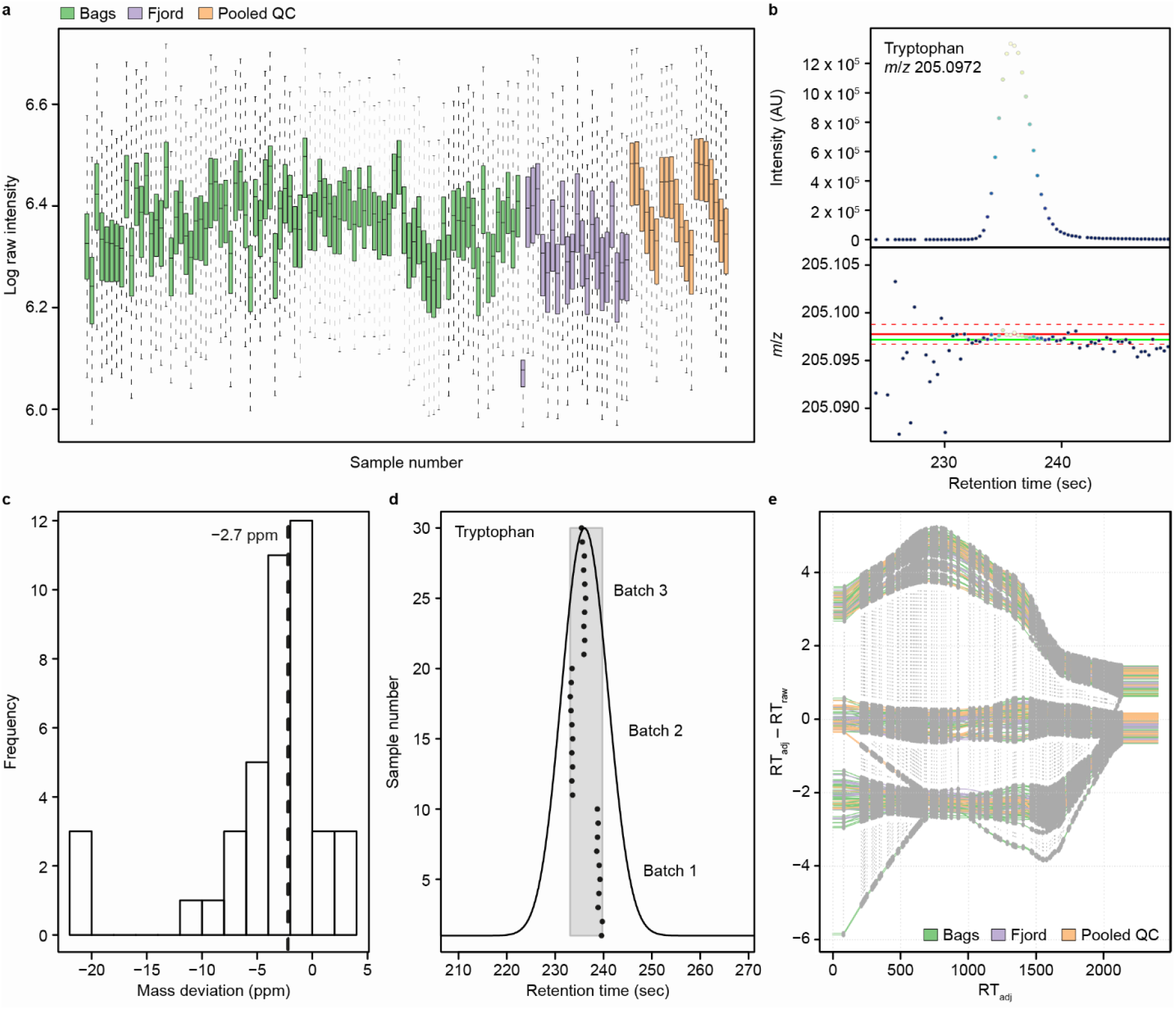
Quality control plots generated during metabolomics data pre-processing. **a**, Log-transformed raw intensity distribution of UPLC-HRMS scans in individual samples, colored by sample type: mesocosm bags (green), fjord water (purple) and pooled QC (orange). The impact of intra- and inter-batch ionization efficiency variances is clearly visible in the pooled QC boxplots. One irregular sample (Fjord, day 10) was detected by having a lower log TIC sum and was marked for removal. **b**, Representative chromatographic peak of an authentic standard (tryptophan, theoretical *m*/*z* 205.0972), which was used to evaluate chromatographic and mass measurement performance (top and bottom panel, respectively). In total, 13 authentic standards were injected as technical repeats in consistent intervals over the whole experiment and inspected for peak quality and parameter adjustments (‘MExMix’; Supplementary Table 4). Red line in bottom panel – theoretical *m*/*z*; dashed red lines – ±5.0 ppm window; green line – measured *m*/*z*. **c**, A histogram of mass measurement errors of the 13 authentic standards measured in the positive ionization mode (n = 3 injections) indicating an overall high accuracy at <3 ppm, with a lower precision of ±12.5 ppm at the 95% confidence level. **d**, Grouping of the tryptophan mass peaks over 30 technical repeats and three analytical batches demonstrates the inter-batch variability in the chromatographic domain (each injection is denoted as a black dot and elutes between 233 and 240 sec). The grey rectangle demonstrates the ability of the grouping algorithm to overcome the variability and identify the fluctuating peaks as a unique group of signals corresponding with one mass feature. Optimized values of three parameters, which highly affect the performance of the grouping algorithm, are: ‘bw’ = 4, ‘binSize’ = 0.05, ‘minFrac’ = 0.7 (Supplementary Table 6). **e**, Retention time (RT) alignment correction curves show a clear inter-batch chromatographic separation of roughly ±6 sec, which corresponds with a stable chromatographic system. Grey dots connected by vertical dotted lines denote groups of consensus peaks, which the alignment algorithm uses to calculate the per-sample retention time correction curves.

**Extended Data Fig. 8:**
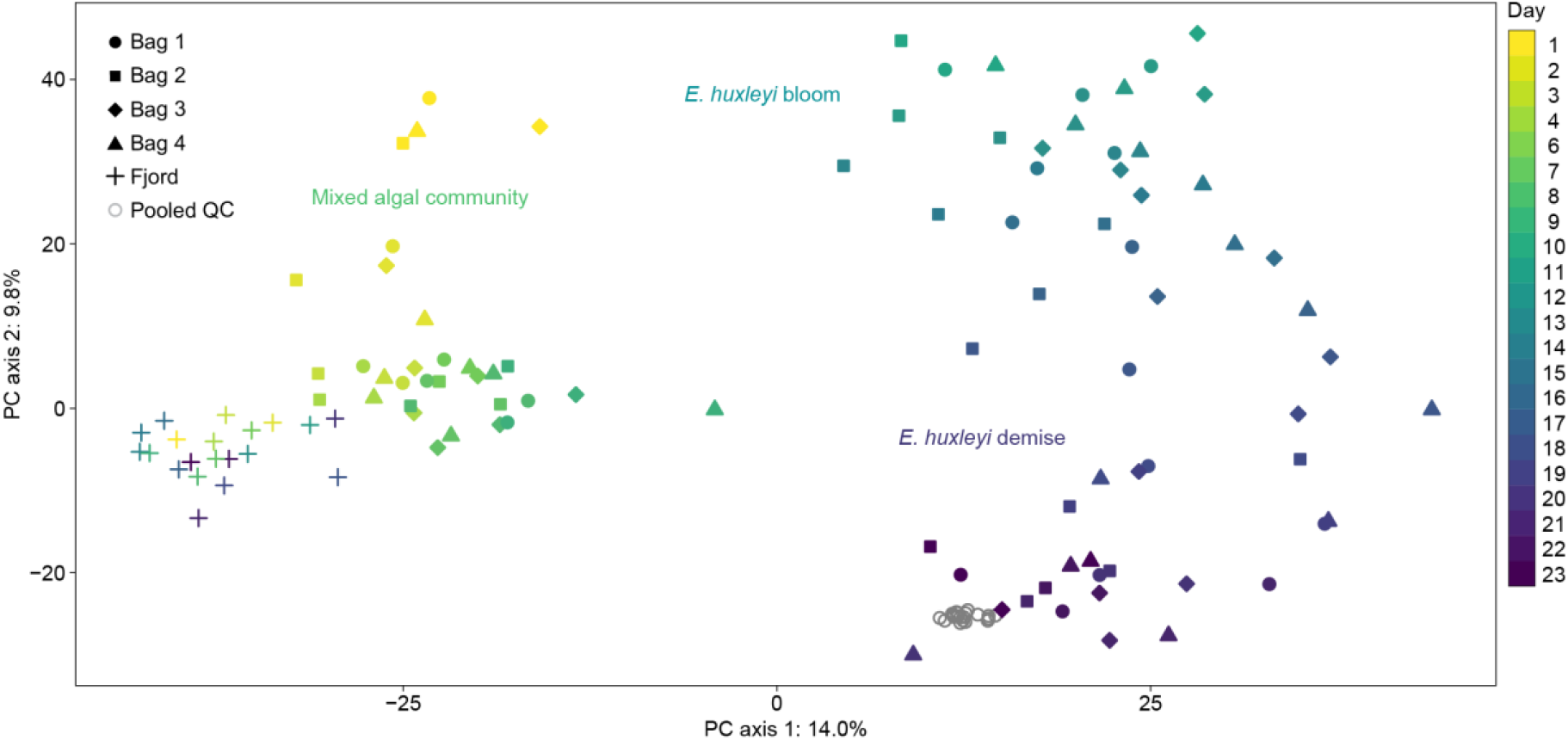
PCA score plot of the exometabolite profiling data acquired in negative ionization mode. Principal component analysis (PCA) of the extracted DOM (based on 4,517 mass features) separates the fjord samples (crosses) from mesocosm bag samples (filled symbols). The day of the experiment is indicated by the color of the symbols, from yellow (day 1) to blue (day 23). PC axis 1 (14.0%) and PC axis 2 (9.8%) reflect phytoplankton bloom succession through time: the fjord and the mixed algal community phase (day 1-9) are separated from the *E. huxleyi* bloom and demise phases (day 10-23) along the first axis. The second axis further separates daily changes within each phase and between the bloom and the demise phase of *E. huxleyi*. Phytoplankton bloom phases are indicated. Pooled QC samples (hollow circles) are shown as a reference.

**Extended Data Table 1:** Exometabolites characterizing the virus-induced DOM of *E. huxleyi* blooms.

| # | Halogenation pattern | RT (min) | Measured *m/z* | Molecular ion | Predicted formula* | Predicted mass | Pred. DBE | Theoretical *m/z* (pos) | Δppm of *m/z* | Possible formula difference<sup>†</sup> |
| --- | --- | --- | --- | --- | --- | --- | --- | --- | --- | --- |
| 1 | Chloro (Cl) | 11.13 | 364.9544 |  |  |  |  |  |  |  |
| 2 | - | 11.42 | 211.0965 | [M+H]<sup>+</sup> | C<sub>11</sub>H<sub>14</sub>O<sub>4</sub> | 210.0865 | 4.5 | 211.0965 | 0.0 |  |
| 3 | Trichloro-Iodo (Cl<sub>3</sub> I) | 13.75 | 639.0027 | [M+H]<sup>+</sup> | C<sub>18</sub>H<sub>30</sub>Cl<sub>3</sub>IO<sub>10</sub> | 637.9949 | 1.5 | 639.0022 | 0.8 | +O |
|  |  |  | 636.9901 | [M-H]<sup>-</sup> |  |  |  |  |  |  |
| 4 | Dichloro-Iodo (Cl<sub>2</sub> I) | 15.15 | 629.0405 | [M+H-H<sub>2</sub>O]<sup>+</sup> | C<sub>20</sub>H<sub>33</sub>Cl<sub>2</sub>IO<sub>11</sub> | 646.0433 | 3.5 | 629.0412 | -1.1 | +C<sub>2</sub>H<sub>2</sub>O<sub>2</sub> |
|  |  | ‡ | 645.0341 | [M-H]<sup>-</sup> |  |  |  |  |  |  |
| **5** | **Dichloro-Iodo (Cl<sub>2</sub> I)** | **15.15** | **589.0464** | **[M+H]<sup>+</sup>** | **C<sub>18</sub>H<sub>31</sub>Cl<sub>2</sub>IO<sub>9</sub>** | **588.0386** | **1.5** | **589.0463** | **0.2** | **Reference** |
|  |  | ‡ | **587.0295** | **[M-H]<sup>-</sup>** |  |  |  |  |  |  |
| 6 | Dichloro-Iodo (Cl<sub>2</sub> I) | 15.55 | 629.0402 | [M+H-H<sub>2</sub>O]<sup>+</sup> | C<sub>20</sub>H<sub>33</sub>Cl<sub>2</sub>IO<sub>11</sub> | 646.0429 | 3.5 | 629.0412 | -1.6 | +C<sub>2</sub>H<sub>2</sub>O<sub>2</sub> |
|  |  |  | 645.0341 | [M-H]<sup>-</sup> |  |  |  |  |  |  |
| 7 | Trichloro-Iodo (Cl<sub>3</sub> I) | 16.90 | 623.0067 | [M+H]<sup>+</sup> | C<sub>18</sub>H<sub>30</sub>Cl<sub>3</sub>IO<sub>9</sub> | 621.9989 | 1.5 | 623.0073 | -1.0 |  |
|  |  |  | 620.9957 | [M-H]<sup>-</sup> |  |  |  |  |  |  |
| 8 | Dichloro-Iodo (Cl<sub>2</sub> I) | 17.13 | 603.0619 | [M+H]<sup>+</sup> | C<sub>19</sub>H<sub>33</sub>Cl<sub>2</sub>IO<sub>9</sub> | 602.0541 | 1.5 | 603.0619 | 0.0 | +CH<sub>2</sub> |
|  |  |  | 601.0474 | [M-H]<sup>-</sup> |  |  |  |  |  |  |
| 9 | Iodo (I) | 17.23 | 618.1921 | [M+H]<sup>+</sup> | C<sub>27</sub>H<sub>40</sub>IO<sub>7</sub>N | 617.1843 | 7.5 | 618.1922 | -0.2 |  |
|  |  |  | 616.1785 | [M-H]<sup>-</sup> |  |  |  |  |  |  |
| 10 | Dichloro-Iodo (Cl<sub>2</sub> I) | 17.25 | 629.0409 | [M+H-H<sub>2</sub>O]<sup>+</sup> | C<sub>20</sub>H<sub>33</sub>Cl<sub>2</sub>IO<sub>11</sub> | 646.0436 | 3.5 | 629.0412 | -0.5 | +C<sub>2</sub>H<sub>2</sub>O<sub>2</sub> |
|  |  |  | 645.0382 | [M-H]<sup>-</sup> |  |  |  |  |  |  |
| 11 | - | 17.70 | 193.0859 | [M+H]<sup>+</sup> | C<sub>11</sub>H<sub>12</sub>O<sub>3</sub> | 192.0786 | 5.5 | 193.0859 | 0.0 |  |
| 12 | Trichloro (Cl<sub>3</sub>) | 21.00 | 366.1149 | [M+H]<sup>+</sup> |  |  |  |  |  |  |
| 13 | Iodo (I) | 24.05 | 600.1817 | [M+H]<sup>+</sup> | C<sub>27</sub>H<sub>38</sub>IO<sub>6</sub>N | 599.1739 | 8.5 | 600.1817 | 0.0 |  |
| 14 | Trichloro-Iodo (Cl<sub>3</sub> I) | 25.08 | 586.9858 | [M+H-H<sub>2</sub>O]<sup>+</sup> | C<sub>18</sub>H<sub>28</sub>Cl<sub>3</sub>IO<sub>8</sub> | 603.9886 | 3.5 | 586.9862 | -0.7 | -H<sub>2</sub>O |
|  |  |  | 602.9799 | [M-H]<sup>-</sup> |  |  |  |  |  |  |
| 15 | - | 25.27 | 363.4955 |  |  |  |  |  |  |  |
| 16 | Dichloro-Iodo (Cl<sub>2</sub> I) | 25.28 | 611.0307 | [M+H-H<sub>2</sub>O]<sup>+</sup> | C<sub>20</sub>H<sub>31</sub>Cl<sub>2</sub>IO<sub>10</sub> | 628.0335 | 4.5 | 611.0306 | 0.2 | +C<sub>2</sub>H<sub>2</sub>O<sub>2</sub> -H<sub>2</sub>O |
|  |  |  | 627.0261 | [M-H]<sup>-</sup> |  |  |  |  |  |  |
| 17 | Trichloro (Cl<sub>3</sub>) | 29.95 | 861.2982 |  |  |  |  |  |  |  |
| 18 | Trichloro (Cl<sub>3</sub>) | 29.98 | 803.2923 | [M+H]<sup>+</sup> |  | 802.2856 |  |  |  |  |
|  |  |  | 801.2789 | [M-H]<sup>-</sup> |  |  |  |  |  |  |
| 19 | Bromo-Chloro (Br<sub>x</sub> Cl<sub>y</sub>) | 30.37 | 905.2472 | [M+H]<sup>+</sup> |  | 904.2382 |  |  |  |  |
|  |  |  | 903.2291 | [M-H]<sup>-</sup> |  |  |  |  |  |  |
| 20 | Bromo-Chloro (Br<sub>x</sub> Cl<sub>y</sub>) | 30.40 | 847.2414 |  |  |  |  |  |  |  |

\*Chemical formulas were predicted based on isotopic pattern and fragmentation tree analysis (see Methods section). †Spectral data indicated a shared core structure for all predicted formulas consisting solely of CHO. The differences between these non-halogenated core structures were thus evaluated. First, a predicted non-halogenated mass was calculated by replacing all halogen atoms with hydrogen atoms. Then, mass differences were calculated towards the most intense compound #5 (in bold). For each mass difference, the molecular formula difference was predicted (<3 mDa). ‡ The co-eluting metabolites #4 and #5 were verified using additional chromatography and MS/MS analysis. DBE = double bond equivalent. For detailed mass spectral information of all MS1 and MS/MS analyses refer to Supplementary Data.

## Supplementary Information Supplementary Methods

### Enumeration of large virus-like particles (VLP) and bacteria using flow cytometry

Extracellular large VLP and bacteria were quantified as described previously^21,57^. Briefly, water samples were fixed with a final concentration 0.5% glutaraldehyde for 30 min at 4°C, plunged into liquid nitrogen and then thawed. 10 µL of fixed sample were stained with SYBR gold (Invitrogen, Paisley, UK) prepared in Tris-EDTA buffer as instructed by the manufacturer (5 μL SYBR gold in 50 mL 0.22 µm filtered Tris-EDTA), then incubated for 20 min at 80°C and cooled down to room temperature. Flow cytometry analysis was performed using an Eclipse iCyt (Sony Biotechology) flow cytometer with excitation at 488 nm and emission at 525 nm. Gates for bacteria and VLPs were set by plotting the emission at 525 nm against the emission at 663-737 nm. A total volume of 30 µL with a flow rate of 10 µL/min was analyzed. A threshold was applied based on the forward scatter signal to reduce the background noise. The gates for large VLP and bacteria were set by comparing to reference samples containing fixed EhV201 and bacteria from lab cultures.

---

**Supplementary Table 1:** Blank and QC samples collected during the mesocosm experiment.

| Name | Type | Amount | Purpose | Description |
| --- | --- | --- | --- | --- |
| Processing blank | Blank | 2 | Detection of contaminations introduced by sample processing, including filtration, extraction and elution | 1 L artificial seawater<sup>58</sup> was processed as a biological sample, as described in the Methods section |
| Extraction blank | Blank | 4<sup>†</sup> | Detection of contaminations introduced by the SPE cartridges | SPE cartridge was processed without loading of a biological sample, as described in the Methods section |
| Pipette blank | Blank | 3 | Detection of contaminations introduced by glassware used during extraction procedure | 4 mL methanol were added using a glass serological pipette to a glass vial |
| Methanol blank | Blank | 4<sup>††</sup> | Detection of contaminations introduced by the elution solvent | 4 mL methanol were added directly to a glass vial |
| IS matrix | IS QC | 2 | Assessing the matrix effect on the ionization efficiency of extraction standards during UPLC-HRMS | Filtrate of a biological sample was processed without addition of internal standard solutions. The internal standard solutions were added directly to the methanol eluate |
| IS positive | IS QC | 4<sup>††</sup> | Assessing the matrix effect on the ionization efficiency of extraction standards during UPLC-HRMS | 4 mL methanol containing internal standards were added directly to a glass vial |

IS – internal standard; QC – quality control. <sup>†</sup>Three out of four samples were injected. <sup>††</sup>Two out of four samples were injected.

**Supplementary Table 2:** Distribution of randomized biological samples, blank samples and IS QC samples between three analytical batches for UPLC-HRMS analysis.

| **Sample type** | **Batch 1** | **Batch 2** | **Batch 3** |
| --- | --- | --- | --- |
| Bag 1 | 9 | 4 | 9 |
| Bag 2 | 9 | 8 | 5 |
| Bag 3 | 6 | 9 | 7 |
| Bag 4 | 6 | 9 | 7 |
| Fjord | 7 | 7 | 8 |
| Blank | 3 | 3 | 4 |
| IS QC | 2 | 2 | 0 |
| **Sum** | **42** | **42** | **40** |

A total of 110 biological samples (22 samples per type) were randomized and divided into three batches, as described in the Methods section. Two blank samples (methanol blank), two IS QC samples (IS positive) and one extraction blank were not used for the analysis.

**Supplementary Table 3:** Sample injection order during UPLC-HRMS analysis.

|  | **Injection number** | **Type** | **Sample** | **Number of injections** |
| --- | --- | --- | --- | --- |
| System | 1 | System suitability blank | Methanol | 1 (neg) |
|  | 2 | System suitability | Authentic standard mixture – Mix15 | 1 (neg) |
|  | 3 | System suitability blank | Methanol | 1 (pos) |
|  | 4 | System suitability | Authentic standard mixture – Mix15 | 1 (pos) |
| Batch (pos/neg) | 5 | System suitability blank | Methanol | 1 |
|  | 6-8 | System suitability | Authentic standard mixture – MExMix | 3 |
|  | 9-11 | Blank | Methanol blank, Extraction blank, Processing blank | 3 |
|  | 12-13 | IS QC | IS matrix, IS positive | 2 |
|  | 14 | Reference Material | EhuxMixM | 1 |
|  | 15-17 | Pooled QC |  | 3 |
|  | 18-27 | Biological sample | Bags, Fjord | 10 |
|  | 28 | Pooled QC |  | 1 |
|  | 29-38 | Biological sample | Bags, Fjord | 10 |
|  | 39 | System suitability | Authentic standard mixture – MExMix | 1 |
|  | 40 | Pooled QC |  | 1 |
|  | 41-50 | Biological sample | Bags, Fjord | 10 |
|  | 51 | Pooled QC |  | 1 |
|  | 52-62 | Biological sample | Bags, Fjord | 7 |
|  | 63 | Pooled QC |  | 1 |
|  | 64 | Reference Material | EhuxMixM | 1 |
|  | 65 | System suitability | Authentic standard mixture – MExMix | 1 |
|  | 66 | System suitability blank | Methanol | 1 |

**Supplementary Table 4:** Chemical composition of authentic standard mixtures and biological reference material.

| Name | Description | Chemical composition |
| --- | --- | --- |
| Mix15 <sup>47</sup> | Authentic standard mixture for long-term comparison of system suitability | Alpha-tomatine (CAS: 17406-45-0)<br>Benzoic acid (CAS: 65-85-0)<br>Caffeic acid (CAS: 331-39-5)<br>Chlorogenic acid (CAS: 327-97-9)<br>Cinnamic acid (CAS: 140-10-3)<br>*p*-Coumaric acid (CAS: 501-98-4)<br>Dehydrotomatine (CAS: 157604-98-3)<br>Ferulic acid (CAS: 537-98-4)<br>Kaempferol (CAS: 520-18-3)<br>(±)-Naringenin (CAS: 67604-48-2)<br>L-Phenylalanine (CAS: 63-91-2)<br>Quercetin (CAS: 117-39-5)<br>Rutin (CAS: 153-18-4)<br>Sinapic acid (CAS: 530-59-6)<br>L-Tryptophan (CAS: 73-22-3)<br>L-Tyrosine (CAS: 60-18-4) |
| MExMix | Authentic standard mixture for intra- and inter-batch comparison of system suitability | Biotin (CAS: 58-85-5)<br>*p*-Coumaric acid (CAS: 501-98-4)<br>2-Heptyl-4-quinolone (CAS: 40522-46-1)<br>C4-homoserine lactone (CAS: 67605-85-0)<br>3-Oxo-C6-homoserine lactone (CAS: 143537-62-6)<br>3-Oxo-C12-homoserine lactone (CAS: 168982-69-2)<br>3-Oxo-C14-homoserine lactone (CAS: 177158-19-9)<br>Indole-3-acetic acid (CAS: 87-51-4)<br>Indole-3-butyric acid (CAS: 133-32-4 )<br>Lumichrome (CAS:1086-80-2)<br>Riboflavin (CAS: 83-88-5)<br>Thymine (CAS: 65-71-4)<br>L-Tryptophan (CAS: 73-22-3) |
| EhuxMixM | Reference material of laboratory cultures for long-term dataset comparison | Solid phase extracts of extracellular metabolites from uninfected and virus-infected cultures of *Emiliana huxleyi* CCMP 2090. Different experimental time points were extracted and all extracts pooled to generate the reference material. |

**Supplementary Table 5:** ‘xcms’ parameters used for peak picking.

| Parameter | Value |
| --- | --- |
| snthresh | 10 |
| ppm | 12 |
| peakwidth_min | 8 |
| peakwidth_max | 18 |
| mzCenterFun | wMeanApex3 |
| noise | 200 |
| prefilter_k | 1 |
| prefilter_I | 250 |

**Supplementary Table 6:** ‘xcms’ parameters used for peak grouping and alignment.

| Parameter | Value |
| --- | --- |
| bw | 4 |
| binSize | 0.05 |
| max | 100 |
| minFraction* | 0.7 |
| smooth | loess |
| family | gaussian |
| span | 0.5 |
| extraPeaks | 0 |

\*For the correlation analysis of metabolite profiling data with extracellular EhV abundance, which used a reduced dataset (only days 10-23), the pre-processing parameter ‘minFraction’ was set to ‘0’ and instead the optional parameter ‘minSamples’ was set to ‘8’, as specified in the Methods section.

**Supplementary Table 7:**
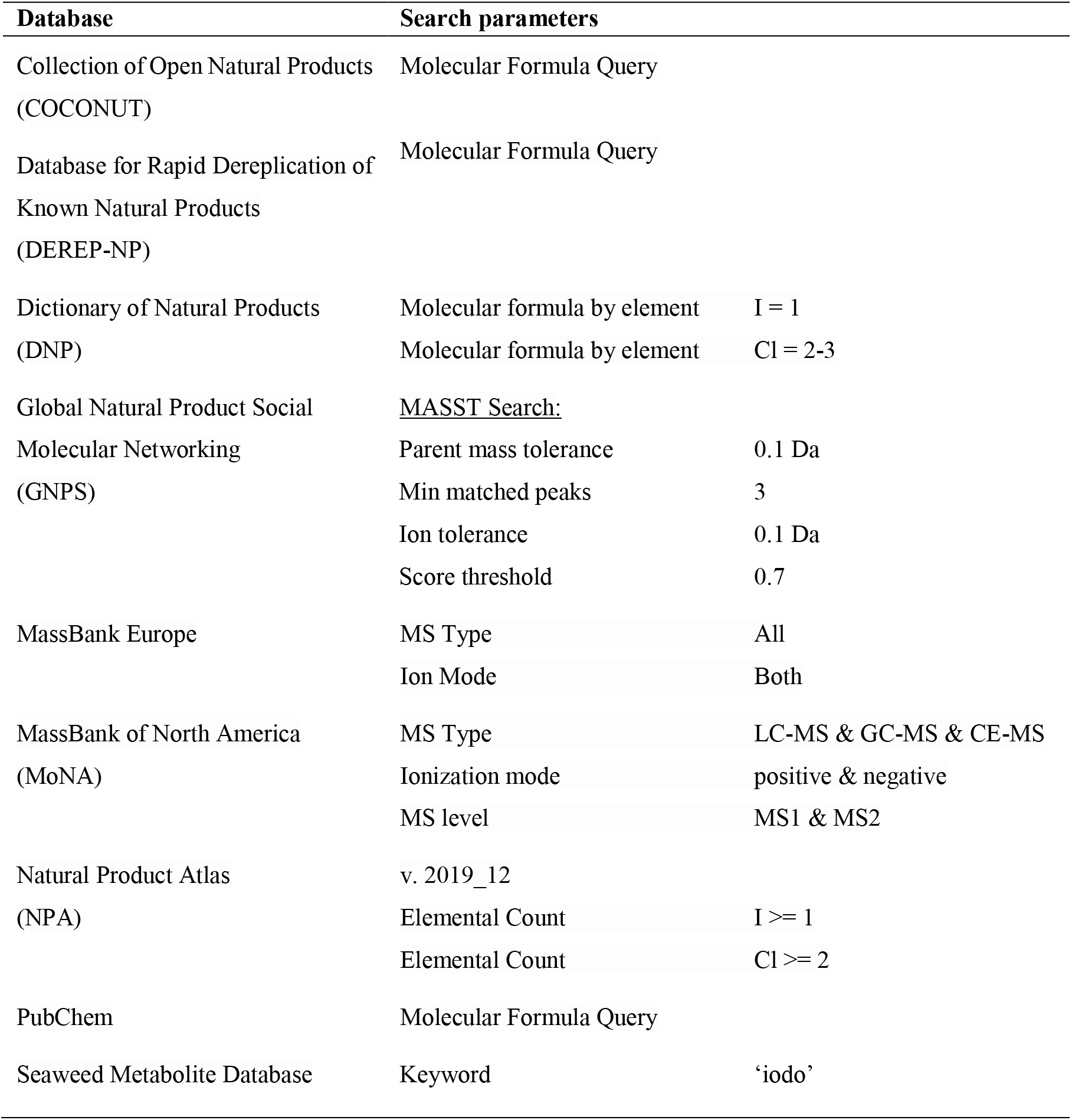
Mass spectral and natural product databases that were queried for the presence of the nine chlorine-iodine-containing metabolites using the predicted molecular formulas C_18_H_28_C_l3_IO_8_, C_18_H_30_Cl_3_IO_9_, C_18_H_30_Cl_3_IO_10_, C_18_H_31_Cl_2_IO_9_, C_19_H_33_Cl_2_IO_9_, C_20_H_31_Cl_2_IO_10_ and C_20_H_33_Cl_2_IO_11_ (representing three isomers).

## References

1. Behrenfeld, M. J. & Boss, E. S. Resurrecting the ecological underpinnings of ocean plankton blooms. Annu. Rev. Mar. Sci. 6, 167–194 (2014).

2 Hansell, D. A., Carlson, C. A., Repeta, D. J. & Schlitzer, R. Dissolved organic matter in the ocean: a controversy stimulates new insights. Oceanography 22, 202–211 (2009).

3 Fuhrman, J. A. Marine viruses and their biogeochemical and ecological effects. Nature 399, 541–548 (1999).

4 Wilhelm, S. W. & Suttle, C. A. Viruses and nutrient cycles in the sea: viruses play critical roles in the structure and function of aquatic food webs. BioScience 49, 781–788 (1999).

5 Zark, M., Christoffers, J. & Dittmar, T. Molecular properties of deep-sea dissolved organic matter are predictable by the central limit theorem: evidence from tandem FT-ICR-MS. Mar. Chem. 191, 9–15 (2017).

6 Hansell, D. A. Recalcitrant dissolved organic carbon fractions. Annu. Rev. Mar. Sci. 5, 421–445 (2013).

7 Biddanda, B. & Benner, R. Carbon, nitrogen, and carbohydrate fluxes during the production of particulate and dissolved organic matter by marine phytoplankton. Limnol. Oceanogr. 42, 506–518 (1997).

8 Kragh, T. & Søndergaard, M. Production and decomposition of new DOC by marine plankton communities: carbohydrates, refractory components and nutrient limitation. Biogeochemistry 96, 177–187 (2009).

9 Thornton, D. C. O. Dissolved organic matter (DOM) release by phytoplankton in the contemporary and future ocean. Eur. J. Phycol. 49, 20–46 (2014).

10 Carlson, C. A. in Biogeochemistry of Marine Dissolved Organic Matter (eds D. A. Hansell & C. A. Carlson) 91–151 (Academic Press, 2002).

11 Ma, X., Coleman, M. L. & Waldbauer, J. R. Distinct molecular signatures in dissolved organic matter produced by viral lysis of marine cyanobacteria. Environ. Microbiol. 20, 3001–3011 (2018).

12 Bratbak, G., Egge, J. K. & Heldal, M. Viral mortality of the marine alga *Emiliania huxleyi* (Haptophyceae) and termination of algal blooms. Mar. Ecol. Prog. Ser. 93, 39–48 (1993).

13 Bratbak, G., Wilson, W. & Heldal, M. Viral control of *Emiliania huxleyi* blooms? J. Mar. Syst. 9, 75–81 (1996).

14 Lehahn, Y. et al. Decoupling physical from biological processes to assess the impact of viruses on a mesoscale algal bloom. Curr. Biol. 24, 2041–2046 (2014).

15 Westbroek, P., Young, J. R. & Linschooten, K. Coccolith production (biomineralization) in the marine alga *Emiliania huxleyi*. J. Protozool. 36, 368–373 (1989).

16 Holligan, P. M. et al. A biogeochemical study of the coccolithophore, *Emiliania huxleyi*, in the North Atlantic. Global Biogeochem. Cy. 7, 879–900 (1993).

17 Brown, C. W. & Yoder, J. A. Coccolithophorid blooms in the global ocean. J. Geophys. Res. Oceans 99, 7467–7482 (1994).

18 Harris, R. P. Zooplankton grazing on the coccolithophore *Emiliania huxleyi* and its role in inorganic carbon flux. Mar. Biol. 119, 431–439 (1994).

19 Stefels, J., Steinke, M., Turner, S., Malin, G. & Belviso, S. Environmental constraints on the production and removal of the climatically active gas dimethylsulphide (DMS) and implications for ecosystem modelling. Biogeochemistry 83, 245–275 (2007).

20 Malitsky, S. et al. Viral infection of the marine alga *Emiliania huxleyi* triggers lipidome remodeling and induces the production of highly saturated triacylglycerol. New Phytol. 210, 88–96 (2016).

21 Schleyer, G. et al. In plaque-mass spectrometry imaging of a bloom-forming alga during viral infection reveals a metabolic shift towards odd-chain fatty acid lipids. Nat. Microbiol. 4, 527–538 (2019).

22 Rosenwasser, S. et al. Rewiring host lipid metabolism by large viruses determines the fate of *Emiliania huxleyi*, a bloom-forming alga in the ocean. Plant Cell 26, 2689–2707 (2014).

23 Vardi, A. et al. Viral glycosphingolipids induce lytic infection and cell death in marine phytoplankton. Science 326, 861–865 (2009).

24 Egge, J. K. & Heimdal, B. R. Blooms of phytoplankton including *Emiliania huxleyi* (Haptophyta). Effects of nutrient supply in different N : P ratios. Sarsia 79, 333–348 (1994).

25 Zark, M., Riebesell, U. & Dittmar, T. Effects of ocean acidification on marine dissolved organic matter are not detectable over the succession of phytoplankton blooms. Sci. Adv. 1, e1500531 (2015).

26 Landa, M., Blain, S., Christaki, U., Monchy, S. & Obernosterer, I. Shifts in bacterial community composition associated with increased carbon cycling in a mosaic of phytoplankton blooms. ISME J. 10, 39–50 (2016).

27 Zimmerman, A. E. et al. Metabolic and biogeochemical consequences of viral infection in aquatic ecosystems. Nat. Rev. Microbiol. 18, 21–34 (2020).

28 Rosenwasser, S., Ziv, C., Creveld, S. G. v. & Vardi, A. Virocell metabolism: metabolic innovations during host-virus interactions in the ocean. Trends Microbiol. 24, 821–832 (2016).

29 Laber, C. P. et al. Coccolithovirus facilitation of carbon export in the North Atlantic. Nat. Microbiol. 3, 537–547 (2018).

30 Butler, A. & Walker, J. V. Marine haloperoxidases. Chem. Rev. 93, 1937–1944 (1993).

31 Lavoie, S. et al. Iodinated meroditerpenes from a red alga *Callophycus* sp. J. Org. Chem. 82, 4160–4169 (2017).

32 Borrelli, F., Campagnuolo, C., Capasso, R., Fattorusso, E. & Taglialatela-Scafati, O. Iodinated indole alkaloids from *Plakortis simplex* − new plakohypaphorines and an evaluation of their antihistamine activity. Eur. J. Org. Chem. 2004, 3227–3232 (2004).

33 Sun, J.-F. et al. Dichotellides A–E, five new iodine-containing briarane type diterpenoids from *Dichotella gemmacea*. Tetrahedron 67, 1245–1250 (2011).

34 Gribble, G. W. Biological activity of recently discovered halogenated marine natural products. Mar. Drugs 13, 4044–4136 (2015).

35 Winter, J. M. & Moore, B. S. Exploring the chemistry and biology of vanadium-dependent haloperoxidases. J. Biol. Chem. 284, 18577–18581 (2009).

36 Sheyn, U., Rosenwasser, S., Ben-Dor, S., Porat, Z. & Vardi, A. Modulation of host ROS metabolism is essential for viral infection of a bloom-forming coccolithophore in the ocean. ISME J. 10, 1742–1754 (2016).

37 Gkotsi, D. S. et al. A marine viral halogenase that iodinates diverse substrates. Nat. Chem. 11, 1091–1097 (2019).

38 Brum, J. R. et al. Patterns and ecological drivers of ocean viral communities. Science 348, 1261498 (2015).

39 Jiao, N. et al. Microbial production of recalcitrant dissolved organic matter: long-term carbon storage in the global ocean. Nat. Rev. Microbiol. 8, 593–599 (2010).

## Method references

40 Vardi, A. et al. Host-virus dynamics and subcellular controls of cell fate in a natural coccolithophore population. Proc. Natl. Acad. Sci. U.S.A. 109, 19327–19332 (2012).

41 Vincent, F. et al. AQUACOSM VIMS-Ehux – Core data. Dryad https://doi.org/10.5061/dryad.q573n5tfr (2020).

42 Sheyn, U. et al. Expression profiling of host and virus during a coccolithophore bloom provides insights into the role of viral infection in promoting carbon export. ISME J. 12, 704–713 (2018).

43 Pagarete, A., Allen, M. J., Wilson, W. H., Kimmance, S. A. & De Vargas, C. Host– virus shift of the sphingolipid pathway along an *Emiliania huxleyi* bloom: survival of the fattest. Environ. Microbiol. 11, 2840–2848 (2009).

44 Everroad, R. C. et al. Concentration of metabolites from low-density planktonic communities for environmental metabolomics using nuclear magnetic resonance spectroscopy. JoVE - J. Vis. Exp., e3163 (2012).

45 Dittmar, T., Koch, B., Hertkorn, N. & Kattner, G. A simple and efficient method for the solid-phase extraction of dissolved organic matter (SPE-DOM) from seawater. Limnol. Oceanogr.-Meth. 6, 230–235 (2008).

46 Team, R. C. R: A language and environment for statistical computing. R Foundation for Statistical Computing, Vienna, Austria. https://www.r-project.org (2020).

47 Shahaf, N. et al. The WEIZMASS spectral library for high-confidence metabolite identification. Nat. Commun. 7, 12423 (2016).

48 Chambers, M. C. et al. A cross-platform toolkit for mass spectrometry and proteomics. Nat. Biotechnol. 30, 918–920 (2012).

49 Smith, C. A., Want, E. J., O’Maille, G., Abagyan, R. & Siuzdak, G. XCMS : processing mass spectrometry data for metabolite profiling using nonlinear peak alignment, matching, and identification. Anal. Chem. 78, 779–787 (2006).

50 Kuhl, C., Tautenhahn, R., Böttcher, C., Larson, T. R. & Neumann, S. CAMERA: an integrated strategy for compound spectra extraction and annotation of liquid chromatography/mass spectrometry data sets. Anal. Chem. 84, 283–289 (2012).

51 van der Kloet, F. M., Bobeldijk, I., Verheij, E. R. & Jellema, R. H. Analytical error reduction using single point calibration for accurate and precise metabolomic phenotyping. J. Proteome Res. 8, 5132–5141 (2009).

52 Dieterle, F., Ross, A., Schlotterbeck, G. & Senn, H. Probabilistic quotient normalization as robust method to account for dilution of complex biological mixtures. Application in ^1^H NMR metabonomics. Anal. Chem. 78, 4281–4290 (2006).

53 Böcker, S. & Dührkop, K. Fragmentation trees reloaded. J. Cheminformatics 8, 5 (2016).

54 Zani, C. L. & Carroll, A. R. Database for rapid dereplication of known natural products using data from MS and fast NMR experiments. J. Nat. Prod. 80, 1758–1766 (2017).

55 Sorokina, M. & Steinbeck, C. Review on natural products databases: where to find data in 2020. J. Cheminformatics 12, 20 (2020).

56 Schieler, B. M. et al. Nitric oxide production and antioxidant function during viral infection of the coccolithophore *Emiliania huxleyi*. ISME J. 13, 1019–1031 (2019).

## References

57 Barak-Gavish, N. et al. Bacterial virulence against an oceanic bloom-forming phytoplankter is mediated by algal DMSP. Sci. Adv. 4(10), eaau5716 (2018).

## References

58 Goyet, C. & Poisson, A. New determination of carbonic acid dissociation constants in seawater as a function of temperature and salinity. Deep Sea Res. Part A. Oceanogr. Res. Pap. 36, 1635-1654 (1989).

